# NECAB2 participates in an endosomal pathway of mitochondrial stress response at striatal synapses

**DOI:** 10.1101/2021.02.15.431234

**Authors:** Diones Bueno, Partha Narayan Dey, Teresa Schacht, Christina Wolf, Verena Wüllner, Elena Morpurgo, Liliana Rojas-Charry, Lena Sessinghaus, Petra Leukel, Clemens Sommer, Konstantin Radyushkin, Michael K.E. Schäfer, Luise Florin, Jan Baumgart, Paul Stamm, Andreas Daiber, Guilherme Horta, Leonardo Nardi, Verica Vasic, Michael J. Schmeisser, Andrea Hellwig, Angela Oskamp, Andreas Bauer, Ruchika Anand, Andreas S. Reichert, Sandra Ritz, Gianluigi Nocera, Claire Jacob, Jonas Peper, Marion Silies, Katrin B. M. Frauenknecht, Axel Methner

## Abstract

Synaptic signaling depends on ATP generated by mitochondria. Due to extensive connectivity, the striatum is especially vulnerable to mitochondrial dysfunction and thus requires efficient mitochondrial quality control and repair. We found that global knockout of the neuronal calcium-binding protein 2 (NECAB2) in the mouse causes loss of striatal synapses and behavioral phenotypes related to striatal dysfunction such as reduced motivation and altered sensory gating. Striatal mitochondria from *Necab2* knockout mice are more abundant and smaller. They are characterized by increased respiration and superoxide production resulting in oxidative stress. This accumulation of dysfunctional mitochondria is caused by a defective assembly of mitochondria with early endosomes in a pathway that involves the small GTPase Rab5 and its guanine nucleotide exchange factor Alsin/ALS2. NECAB2 therefore participates in an endosomal pathway of mitochondrial stress response and repair important for striatal function.

## Introduction

The striatum is an important brain structure involved in motor and action planning, decision-making, motivation and reward (Cox and Witten, 2019). Afferent and efferent projections from the striatum project to other brain regions such as the cortex, thalamus and the substantia nigra (Cox and Witten, 2019). The integrity of this system critically depends on mitochondrial wellbeing because either mitochondrial disease, depletion of mitochondria, or mitochondrial toxins specifically induce striatal degeneration. A point mutation in mitochondrial DNA results in a specific striatal degeneration called Leber’s disease (Novotny et al., 1986), depletion of mitochondrial DNA (Pickrell et al., 2011) also causes striatal degeneration, as does ingestion of the mitochondrial toxin 3-nitropropionic acid (reviewed in (Brouillet et al., 2005)). Furthermore, Huntington’s disease (HD), a hereditary disorder caused by an expanded poly-glutamine in the ubiquitously expressed protein huntingtin, causes striatal degeneration and is associated with mitochondrial dysfunction (reviewed in (Bates et al., 2015)).

It is therefore evident that the striatum depends on the optimal functioning of mitochondria – even more so than other parts of the brain. The reasons are unknown but might be attributed to the high energy demands of the glutamatergic input from both the cortex and thalamus as well as the dopaminergic input from the midbrain. Synaptic signaling consumes a substantial amount of the total ATP generated in the brain. Specifically, glutamatergic synapses have very high energy demands required to reverse the ion movements that generate postsynaptic responses (Harris et al., 2012).

Cells adapt the amount and quality of mitochondria to their metabolic demand by a process called mitochondria quality control which either removes unfit mitochondria via selective mitophagy or oxidized pieces of mitochondria via mitochondria-derived vesicles. Of note, most of these processes involve PTEN-induced putative kinase protein 1 (PINK1) and the E3 ubiquitin ligase Parkin (Clark et al., 2006; Park et al., 2006). In healthy mitochondria, PINK1 is imported and rapidly degraded, but in unfit mitochondria with a reduced mitochondrial membrane potential (ΔΨ_m_), PINK1 import is stalled, leaving enough PINK1 at the outer mitochondrial membrane to phosphorylate Parkin. Parkin then ubiquitinates itself and proteins of the outer mitochondrial membrane (Sarraf et al., 2013) resulting in mitophagy (reviewed by (Narendra et al., 2012)). Importantly, basal mitophagy *in vivo* occurs even in the absence of the PINK1/Parkin pathway in all tissues with high metabolic demand including the brain (McWilliams et al., 2018). Recently, two alternative mitochondrial stress response mechanisms involving Ras-related protein Rab-5 (Rab5)-positive early endosomes were described. These can be triggered either by dissipation of ΔΨ_m_ involving Parkin and the endosomal sorting complexes required for transport (ESCRT) machinery (Hammerling et al., 2017), or by oxidative stress elicited by hydrogen peroxide (H_2_O_2_) mediated by the Rab5 guanine nucleotide exchange factor (GEF) Alsin/ALS2 (Hsu et al., 2018). The relevance of these novel pathways of mitochondrial stress response for striatal function is unclear.

The striatum is the major site of expression of the neuronal Ca^2+^-binding protein 2 (NECAB2), a largely uncharacterized protein containing two EF-hand Ca^2+^-binding domains and a so-called antibiotic biosynthesis monooxygenase (ABM) domain connected by two coiled-coil domains. NECAB2 has two 39 kDa and 43 kDa splice isoforms that differ in 37 amino acids at the N-terminal part of the molecule (Canela et al., 2007). In the striatum, NECAB2 co-localizes and interacts with the G-protein coupled receptors (GPCRs) adenosine A_2A_ receptor (A_2A_R) (Canela et al., 2007) and the metabotropic glutamate receptor 5 (mGluR_5_) (Canela et al., 2009). Functionally, the presence of NECAB2 increases receptor surface localization and basal signal transduction via extracellular signal regulated kinases (ERK) 1 and 2 (Canela et al., 2009, 2007) in a Ca^2+^-dependent manner as the interaction of NECAB2 with either receptor decreases in the presence of high Ca^2+^ levels (Canela et al., 2009, 2007). Ca^2+^ binding is mediated by the EF hands and protein-protein interaction by one of the coiled-coil domains (Canela et al., 2007). The role of the ABM domain in this process is however less obvious. Besides the highly homologous NECAB1 and NECAB3 proteins, ABM domains are only found in a group of atypical heme monooxygenases from bacterial species (reviewed by (Lyles and Eichenbaum, 2018)) and unicellular green algae (Lojek et al., 2017) but not in any other mammalian protein. The substrate of this enzymatic activity and the role of this hypothetical mechanism for striatal function are unclear.

We here studied the role of NECAB2 in striatal brain function and found that global knockout of NECAB2 in the mouse causes loss of striatal synapses and behavioral phenotypes related to striatal dysfunction such as reduced motivation and altered sensory gating. Striatal mitochondria from *Necab2* knockout mice are more abundant but less efficient. They are characterized by increased respiration and superoxide production. This mitochondrial dysfunction is caused by a defective endosomal mitochondrial stress response mediated by the Rab5 GEF Alsin.

## Results

### NECAB2 is expressed in a punctate synaptic and neuronal pattern in the striatum of mice and men

To elucidate the role of NECAB2 in brain function, we first investigated the expression pattern of NECAB2 in the mouse and human brain using immunohistochemistry. This demonstrated that NECAB2 is strongly expressed in the mouse caudate putamen (the striatum), the hippocampal CA2 region, the indusium griseum, habenulae, fasciculus retroflexus, and the substantia nigra (Figure 1A). In the human striatum, NECAB2 is prominently expressed in neurons and the neuropil of the putamen and slightly less in the caudate nucleus (Figure 1B). These two structures constitute the human equivalent of the rodent striatum. In the globus pallidus externus and internus only the neuropil but not the somata of neurons are stained (Figure 1B). The majority of synapses of pallidal neurons represent striatopallidal terminals. Within the striatum, the vast majority of neurons are GABAergic inhibitory medium spiny neurons (MSNs), which are characterized by large and extensive dendritic trees and a high abundance of spines. There are at least two subtypes of MSNs: direct pathway MSNs excite their ultimate basal ganglia output structure and express dopamine D1 receptors and adenosine A1 receptors, while indirect pathway MSNs inhibit their output structure and express dopamine D2 receptors and NECAB2-interacting A_2A_R receptors. Using mice that express tagged dopamine D1 and D2 receptors (Thibault et al., 2013), we found NECAB2 expressed in MSNs of both the direct and indirect pathway (Figure 1C). In mice and men, NECAB2 immunoreactivity is mainly present in intracellular punctate structures located in the somatodendritic and synaptic compartments of medium spiny neurons.

**Figure 1.**
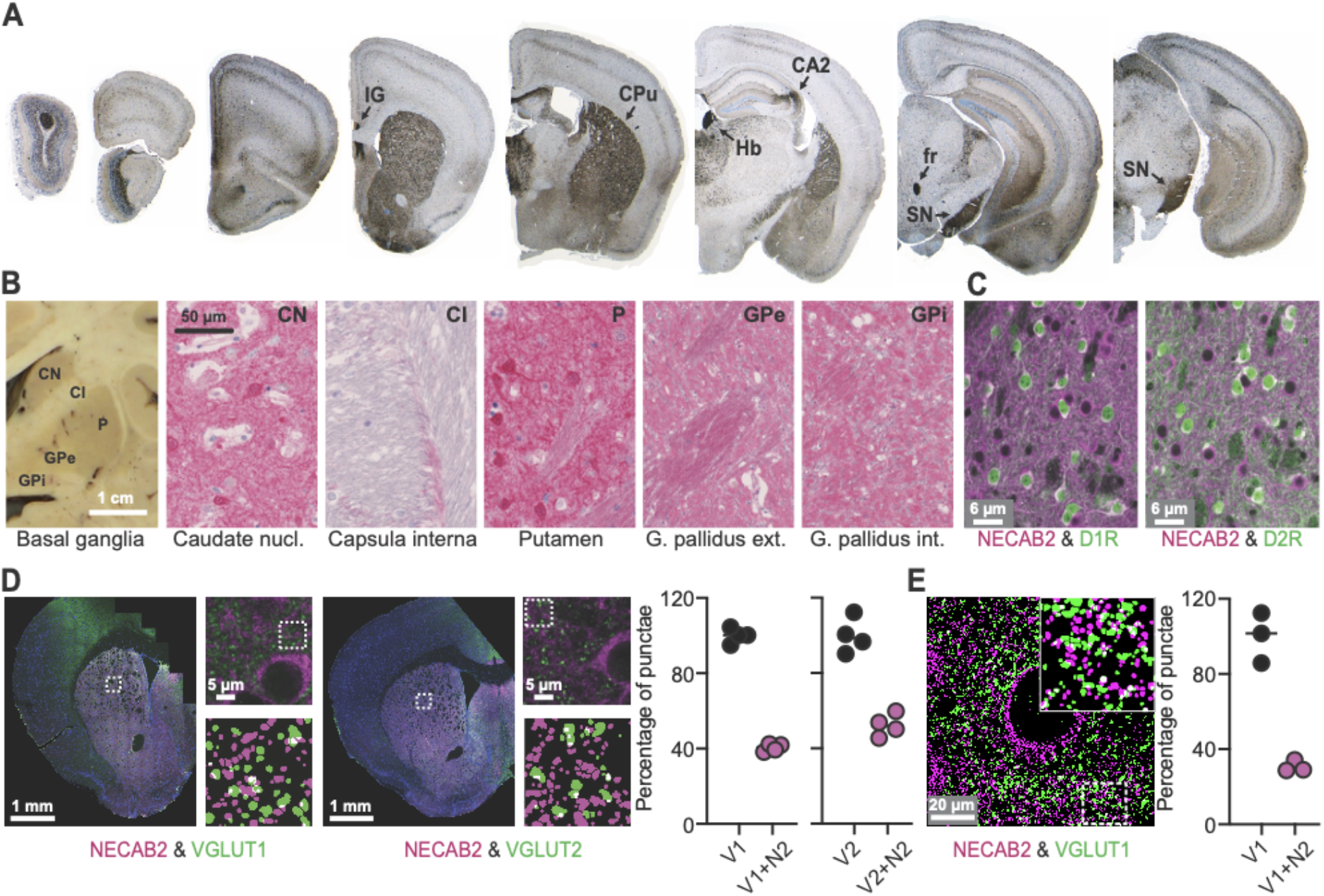
NECAB2 is expressed in a punctate synaptic and neuronal pattern in the striatum of mice and men. (A,B) Immunohistochemical staining demonstrates prominent expression of NECAB2 in (A) the mouse striatum (caudate putamen, CPu), the CA2 region of the hippocampus, indusium griseum (IG), habenulae (Hb), fasciculus retroflexus (fr), and substantia nigra (SN). (B) In the human striatum, which is composed of the caudate nucleus and the putamen, NECAB2 is expressed intracytoplasmically in neurons and in a punctate synaptic pattern in the neuropil. (C) NECAB2 is expressed in medium spiny neurons of the direct (D1R+) and indirect (D2R+) pathway assessed by immunohistochemistry in D1R-tdtomato and D2R-GFP-overexpressing mice. (D) In the mouse striatum, NECAB2 has a punctate staining pattern located in the somatodendritic and synaptic compartments of medium spiny neurons. Partial overlap with both VGLUT1 and VGLUT2 indicates its postsynaptic localization at both corticostriatal and thalamostriatal contacts. Each data point was obtained from one mouse. (E) Expansion microscopy corroborated these results. Size is indicated.

We therefore studied the cellular localization of NECAB2 in relation to synaptic marker proteins that label the presynaptic glutamatergic terminals of corticostriatal (VGLUT1) and thalamostriatal (VGLUT2) synapses. This demonstrated that NECAB2 punctae are in about 40% adjacent to and partially overlapping with VGLUT1 or VGLUT2 signals (Figure 1D), implicating a cytosolic and also partial synaptic localization of NECAB2. These results were reproduced by expansion microscopy which also demonstrated partial colocalization of VGLUT1 and NECAB2 (Figure 1E). Together these results suggest a role for NECAB2 in organelle function and synaptic signaling in the striatum.

### NECAB2 is mainly localized in early endosomes

To clarify the identity of the intracellular NECAB2-immunopositive punctate structures, we used super-resolution STED microscopy of mature primary striatal cultures at day *in vitro* (DIV) 12. Mice with a global knockout (KO) of *Necab2* served as control. Immunoblotting of brain tissue (Figure 2A), immunohistochemistry of brain tissue sections (supplemental Figure 1), and immunocytochemistry of striatal cultures (Figure 2B) demonstrated a complete loss of protein expression in these mice and the specificity of the antibody. Staining wildtype striatal cultures with antibodies against NECAB2 and the endosomal marker Rab5, a small GTPase that is required for the fusion of plasma membranes and early endosomes (reviewed in (Langemeyer et al., 2018), and analysis by STED microscopy revealed an almost complete overlap with a rather neglectable expression of NECAB2 at the plasma membrane (Figure 2C). NECAB2 expression also partially overlapped with CD63, a marker for multivesicular bodies, LC3, a marker for lysosomes, and PDI as a marker for the endoplasmic reticulum (supplemental Figure S2). We also corroborated the predominant endosomal localization of NECAB2 biochemically and found that NECAB2 is present in early endosomal fractions obtained from mouse striatum (Figure 2B) in line with the colocalization with Rab5. We conclude that NECAB2 is mainly found in early endosomes of striatal neurons.

**Figure 2.**
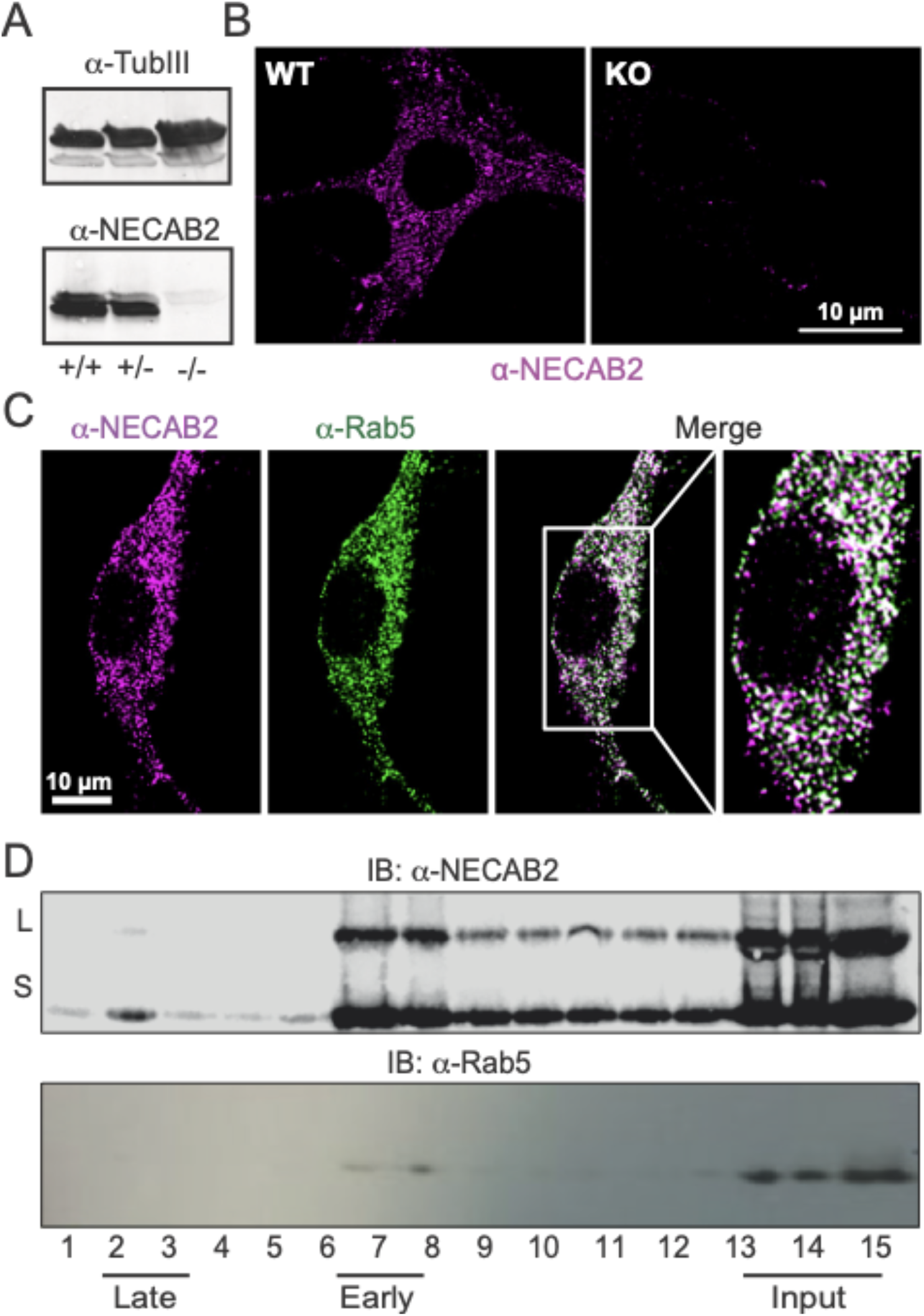
NECAB2 is mainly localized in early endosomes. (A,B) Validation of NECAB2 deficiency and antibody specificity in -/- mice shown by (A) immunoblotting and (B) immunocytochemistry of primary striatal cultures at day in vitro 12. Tubulin-III served as loading control, size is indicated. (C) Punctate pattern of NECAB2 staining in primary striatal cultures colocalizes with the endosome marker Rab5. (D) Presence of NECAB2 in immunoblotted endosomal fractions. L and S represent the long and short splice isoforms of NECAB2. Rab5 served as control.

### NECAB2 deficiency causes striatal dysfunction by loss of striatal synapses

We next asked whether NECAB2 deficiency has consequences for mouse behavior and focused on behavioral functions of the striatum, the major site of NECAB2 expression. In this location, GABAergic striatal MSNs are critical for motor control and play an important role in motivation and reward. We used 15 adult male WT and KO mice and analyzed their behavior using well-established tests. First of all, basic locomotion tested by the open field test or the elevated plus maze test were not impaired (Supplemental Figure S3). However, in line with striatal and basal ganglia dysfunction, we found reduced motivation in the progressive ratio schedule test, increased catalepsy, and reduced sensory gating as analyzed by pre-pulse inhibition of the startle response in KO mice (Figure 3A).

**Figure 3.**
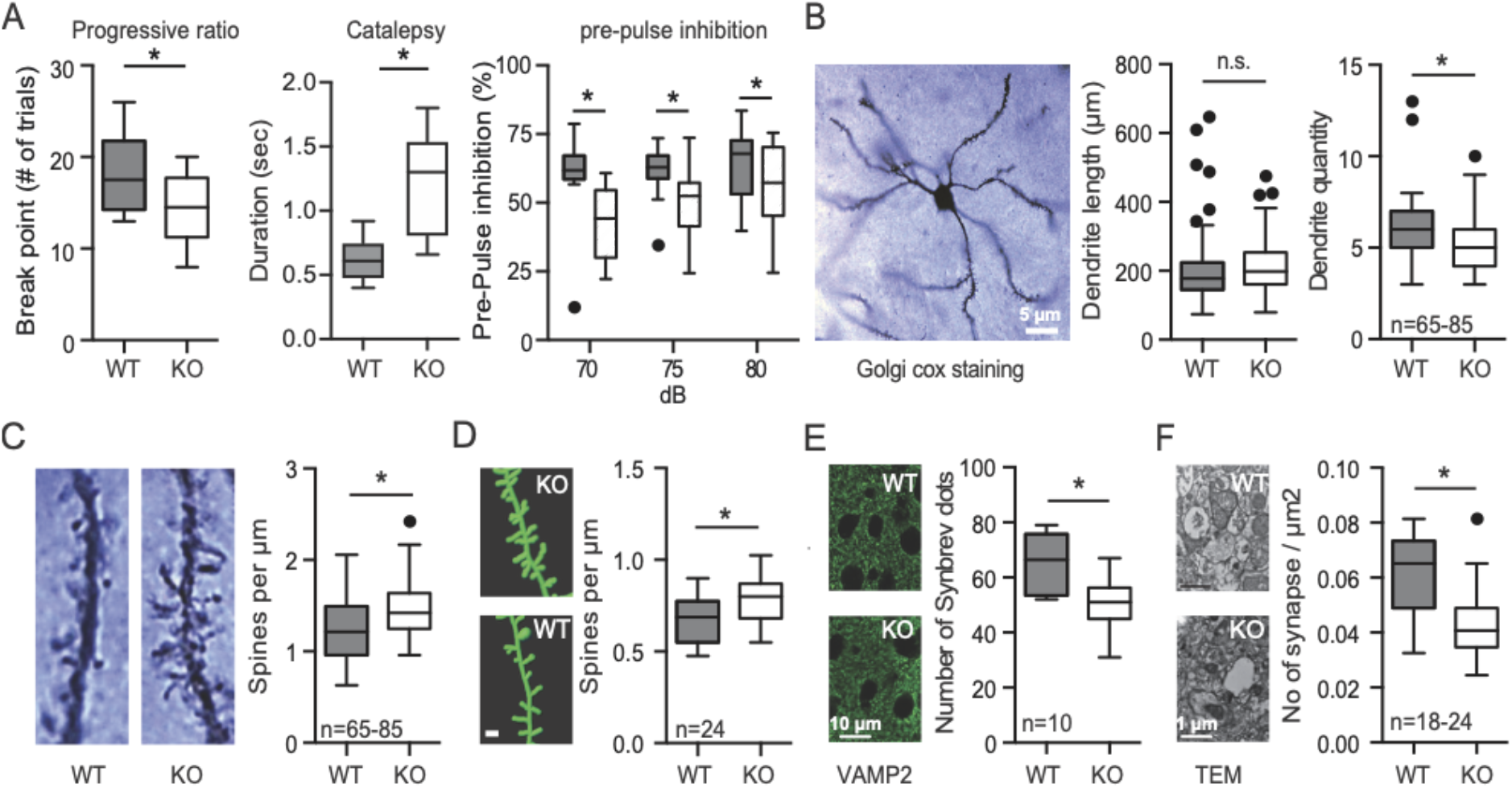
NECAB2 deficiency causes loss of striatal synapses and behavioral deficits associated with striatal dysfunction. (A) Behavioral analysis of 15 male WT and KO adult mice identified a striatal dysfunction with reduced motivation analyzed by the progressive ratio schedule, increased catalepsy upon the i.p. injection of saline, and reduced sensory gating analyzed by pre-pulse inhibition of the startle reflex. (B,C) Golgi cox staining demonstrates (B) a decreased dendrite quantity but (C) more spines per µm. (D) Similar increase in striatal spine density in in-utero electroporated mice expressing GFP. Scale bar 1 µm. (E) Reduced number of striatal synapses in KO mice shown by VAMP2 staining, and (E) transmission electron microscopy. Data are shown as Tukey plots, *<0.05 Mann Whitney test.

Astonishingly, these behavioral changes appear to be independent of alterations of striatal GPCRs, whose function and trafficking is affected by NECAB2 (Canela et al., 2009, 2007). Using autoradiography, we investigated changes in the abundance of the GPCRs interacting with NECAB2 (A_2A_R and mGluR_5_), of receptors known to heterodimerize with these GPCRs (Ferré et al., 2002) that define MSN subsets (D1R and D2R), and of the dopamine transporter (DAT) as a general striatal marker and found no differences in binding density between wildtype and *Necab2* KO mice (Supplemental Figure S4A). Also baseline ERK1 and 2 phosphorylation was similar in wildtype (WT) and KO mice (Supplemental Figure S4B) despite the positive effect of ectopically expressed NECAB2 on A_2A_R (Canela et al., 2007) and mGluR_5_-mediated ERK1/2 phosphorylation in human embryonic kidney cells (Canela et al., 2009). *Necab2* deficiency therefore does not alter the function or abundance of striatal GPCRs – suggesting additional, novel roles for NECAB2.

As a first step to identify such additional roles, we investigated morphological changes associated with this striatal dysfunction by analyzing differences in dendritic arborization and spine density of MSNs using the Golgi-Cox technique. This revealed a similar length, but decreased quantity of dendrites in KO animals (Figure 3B) as well as an increased spine density (Figure 3C). Spine density might serve to compensate for the fewer dendrites. To confirm this finding, we also quantified spines from mice that were *in-utero* electroporated with a vector expressing GFP (Figure 3D). This again demonstrated an increased spine density in the Necab2 KO mouse. In addition, *Necab2* KO mice had a reduced number of striatal synapses shown by immunohistochemistry using the presynaptic marker protein vesicle-associated membrane protein 2 (VAMP2), (Figure 3E). This reduced number of striatal synapses was further confirmed by TEM analysis (Figure 3F). Together these results imply that NECAB2 deficiency causes striatal dysfunction by loss of striatal synapses.

### *Necab2* knockout results in mitochondrial dysfunction

When we studied the number of synapses by electron microscopy, we noted morphological differences of synaptic mitochondria and decided to investigate this in more detail. Because NECAB2 is localized partially at striatal synapses (Figure 1D and E) and affects the number of synapses (Figure 3E/F), we first focused on synaptic mitochondria which support synaptic transmission, a very energy-demanding process. ATP generated by mitochondria is necessary to power ion pumps, support ion gradients, and maintain vesicle recycling and mitochondrial movement (Harris et al., 2012). NECAB2 is indeed abundant in the striatal synaptosomes used for the analysis (Figure 4A). Synaptic *Necab2 KO* mitochondria were significantly smaller and had a higher electron density than those of wildtype samples (Figure 4B). The same mitochondrial morphology and electron density was also evident in tissue sections of the striatum of *Necab2 KO* mice (Figure 4C). Higher electron density is frequently associated with an increased activity of the respiratory chain (Hackenbrock, 1966). Striatal sections of the KO also had an increased number of mitochondria (Figure 4D). Without NECAB2, striatal mitochondria are therefore smaller, more abundant and display a morphological hallmark suggestive of an increased respiratory activity.

**Figure 4.**
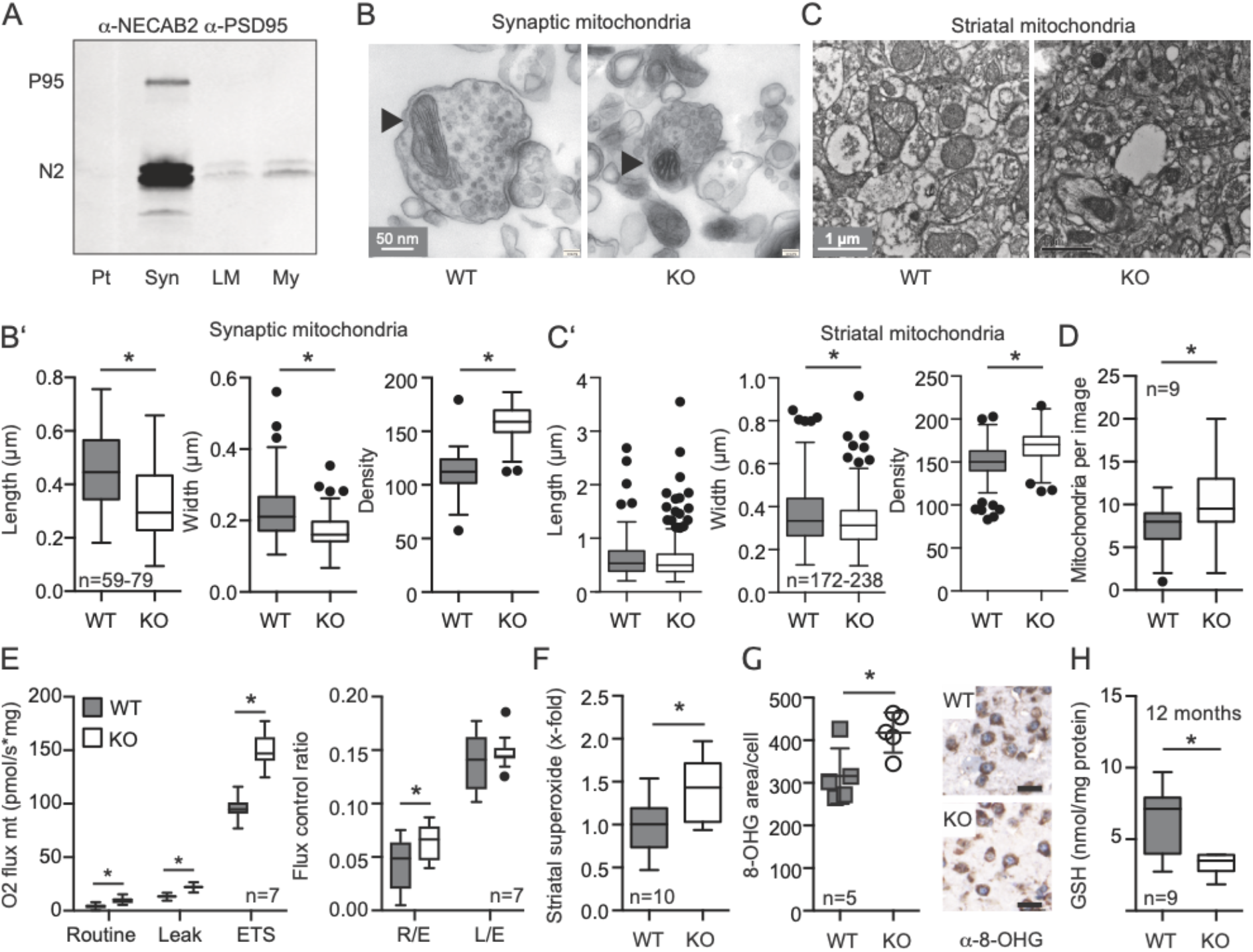
*Necab2* knockout results in mitochondrial dysfunction. **(A)** Immunoblot showing that NECAB2 is enriched in synaptosomes (Syn) but less in pellet (Pt), light membrane (LM) and myelin (My) containing fractions. PSD95 serves as a synaptic marker. **(B**,**C)** Transmission electron microscopy demonstrates morphological alterations of mitochondria in (B) striatal synaptosomes, n=59-79 synapses from 10 mice and in (C) striatal tissue, n=172-238 synapses from 5 mice. KO mitochondria were significantly smaller and showed a higher electron density. **(D)** Increased number of mitochondria in striatal tissue of the KO, quantified per visual field, n=48 from 9 mice. **(E)** Increased respiration of KO mitochondria shown by high-resolution respirometry of striatal tissue. Leak respiration was measured after addition of oligomycin to inhibit the ATP synthase and the electron transfer system (ETS) capacity after titration of FCCP. Flux control ratios are obtained by dividing routine (R) and leak (L) respiration with the ETS (E) values, n=7 mice. **(F)** Increased mitochondrial superoxide production in striatal KO tissue treated with the complex III inhibitor myxothiazol. Superoxide formation was quantitated by the dihydroethidium oxidation product 2-hydroxyethidium detected by its fluorescence at 480/580 nm (Ex/Em) and normalized to wet weight, n=10 mice **(G)** Increased immunohistochemical staining of the reactive oxygen species marker 8-hydroxyguanine (8-OHG) in KO striata. Scale bar represents 20 µm. Each data point contains the mean of n=10 cells analyzed from 5 mice, *<0.05 Mann Whitney test. **(H)** Reduced striatal total glutathione determined enzymatically in adult KO mice, n=6-9 mice. Data are presented as Tukey plots, *<0.05 Mann Whitney test.

We therefore directly investigated mitochondrial respiration in striatal tissue dissected from *Necab2* KO mice. High-resolution respirometry of this striatal tissue demonstrated an increased respiration in the absence of NECAB2 in line with the higher electron density of KO mitochondria. Normalizing respiration to the maximum electron transfer capacity (ETS) revealed that *Necab2* KO mitochondria spend a higher proportion of their maximum capacity for routine respiration and have thus a reduced spare capacity (Figure 4E). This increased ratio between non-phosphorylating leak respiration (electron flow coupled to proton pumping to compensate for proton leaks) and ETS capacity suggests an intrinsic uncoupling or dysfunction of mitochondria in *Necab2* KO tissue. Mitochondria *in situ* also produced more superoxide as assessed by quantifying oxidized 2-hydroxyethidium by high-performance liquid chromatography in striatal tissue treated with the complex III inhibitor myxothiazol (Kröller-Schön et al., 2014) (Figure 4F). We also detected increased amounts of oxidized DNA in striatal tissue using an antibody against 8-hydroxy-2’-deoxyguanosine (Figure 4G) and reduced levels of the major antioxidant glutathione in the striata of adult mice (Figure 4H) in line with a more oxidative environment. Together, these data demonstrate that striatal mitochondria from *Necab2* KO mice are dysfunctional.

### NECAB2 partially colocalizes with mitochondria

To investigate how NECAB2 deficiency could affect mitochondrial wellbeing, we first studied whether NECAB2 also colocalizes with mitochondria and stained striatal tissue sections with antibodies against NECAB2 and the mitochondrial marker protein cytochrome *c* (cyt *c*). This revealed the presence of cyt *c*-positive structures within NECAB2-stained areas (Figure 5A, see also supplemental video 1). Also in primary striatal neurons, NECAB2 partially colocalized with HSPA9, a mitochondrial matrix protein (Rath et al., 2021) (Figure 5B) as demonstrated by superresolution microscopy. Furthermore, NECAB2 is enriched in fractions containing synaptic but not non-synaptic mitochondria obtained from striatal tissue (Figure 5C). Proteinase K digestion of the mitochondrial fraction effaced immunoreactivity against NECAB2 and the voltage-dependent anion channel (VDAC) at the outer mitochondrial membrane but not against the matrix protein cyt *c* oxidase (Figure 5C) indicating that NECAB2 associates with the outer mitochondrial membrane. These results suggest a role for NECAB2 at the interface of mitochondrial and endosomal function primarily at striatal synapses.

**Figure 5.**
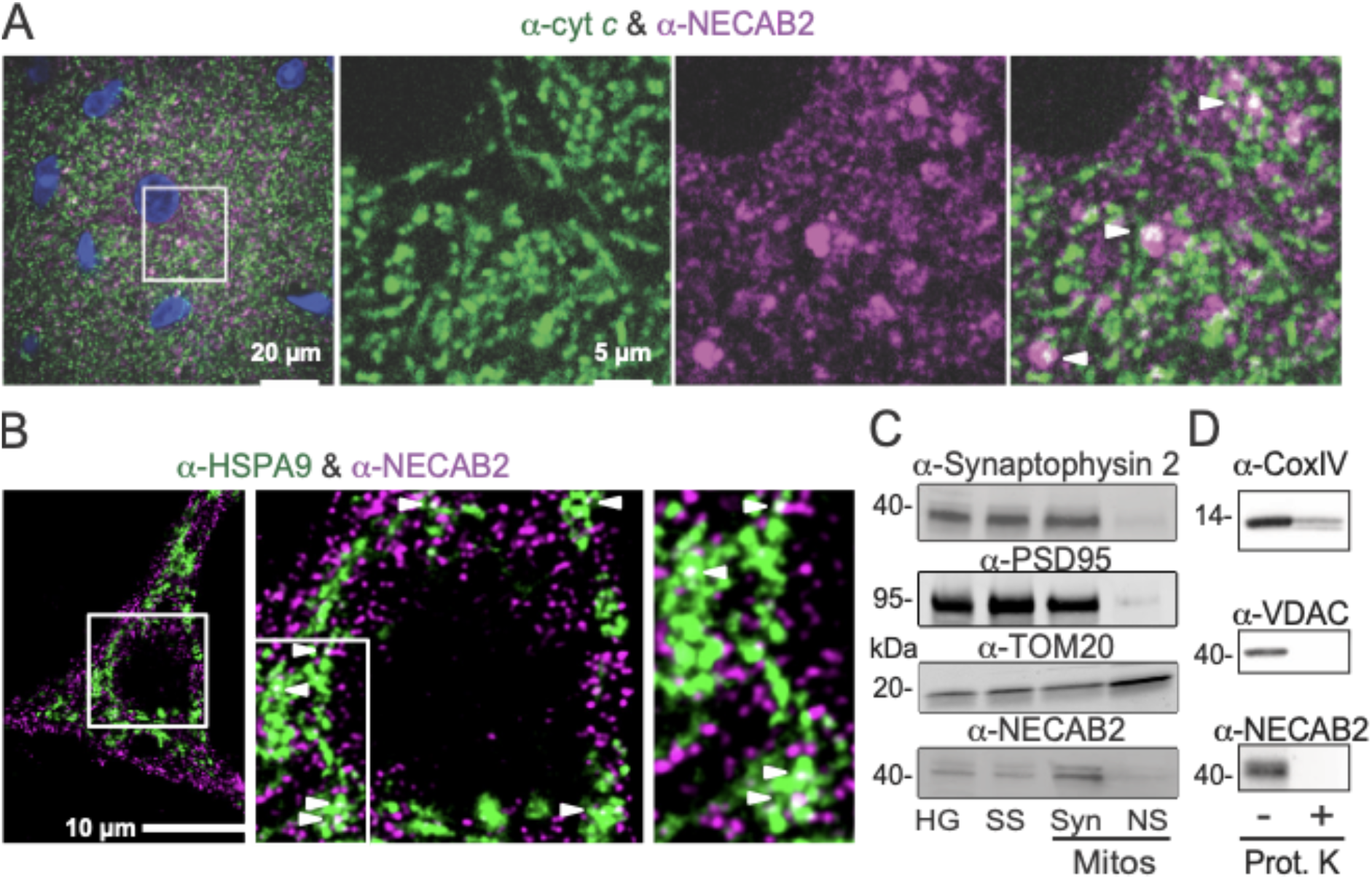
NECAB2 partially co-localizes with mitochondria. (A) Immunohistochemical co-staining of NECAB2 and the mitochondrial protein Cytochrome *c* (cyt *c*) in striatal tissue sections demonstrating that NECAB2-positive punctae contain mitochondrial material. (B) Immunocytochemistry of primary striatal neurons stained with NECAB2 and the mitochondrial marker protein HSPA9. The white arrowheads in A and B point to areas of colocalization. Size is indicated. (C) Presence of NECAB2 in fractions containing synaptic but not non-synaptic mitochondria. PSD95 and synaptophysin-2 served as synaptic markers, TOM20 as mitochondrial marker protein. (D) In mitochondrial fractions, NECAB2 can be digested with proteinase K suggesting its presence at the outer mitochondrial membrane. VDAC as outer mitochondrial membrane protein and cytochrome oxidase IV (CoxIV) as mitochondrial matrix protein served as controls.

### NECAB2 deficiency abrogates endosomal stress-activated mitochondrial signaling elicited by oxidative stress

The preponderance of dysfunctional mitochondria in the absence of NECAB2 (Figure 4) and the association of mitochondrial structures with NECAB2-stained vesicular structures (Figure 5), suggest that mitochondria are less efficiently recycled or repaired by endosomes in the *Necab2* KO. Two recently described and probably distinct mitochondrial stress response pathways involving Rab5-positive early endosomes can be triggered either by dissipation of the mitochondrial membrane potential, involving PINK1/Parkin (Hammerling et al., 2017) or by oxidative stress (Hsu et al., 2018). Increased production of reactive oxygen species (ROS) is a hallmark of dysfunctional mitochondria (Murphy, 2009). To test and distinguish these two pathways, we treated mature primary striatal cultures from WT and KO littermates with FCCP or H_2_O_2_ and quantified endosomes alone, as well as endosomes associated with mitochondria (see exemplary striatal neuron with labeled endosomes and mitochondria in Figure 6A). When comparing just Rab5-positive endosomes, this demonstrated a similar increase in WT and KO neurons upon treatment with FCCP and no changes with H_2_O_2_ (Figure 6B). The abundance of structures positive for both Rab5 and TOM20 (translocase of outer membrane 20, a mitochondrial marker protein) was however different between WT and Necab2 KO. When treated with H_2_O_2_, WT showed a tendency towards an increase in double-positive structures, whereas their number significantly declined in the KO (Figure 6C). In contrast, when treated with FCCP, WT mice tended to show less double-positive structure, which however significantly increased in KO (Figure 6C). This argues that the H_2_O_2_-mediated endosomal stress-activated mitochondrial signaling pathway is compromised in *Necab2* KO animals. The upregulation of double-positive structures in KO upon exposure to FCCP probably represents a compensatory upregulation of the PINK1/Parkin endosomal pathway.

**Figure 6.**
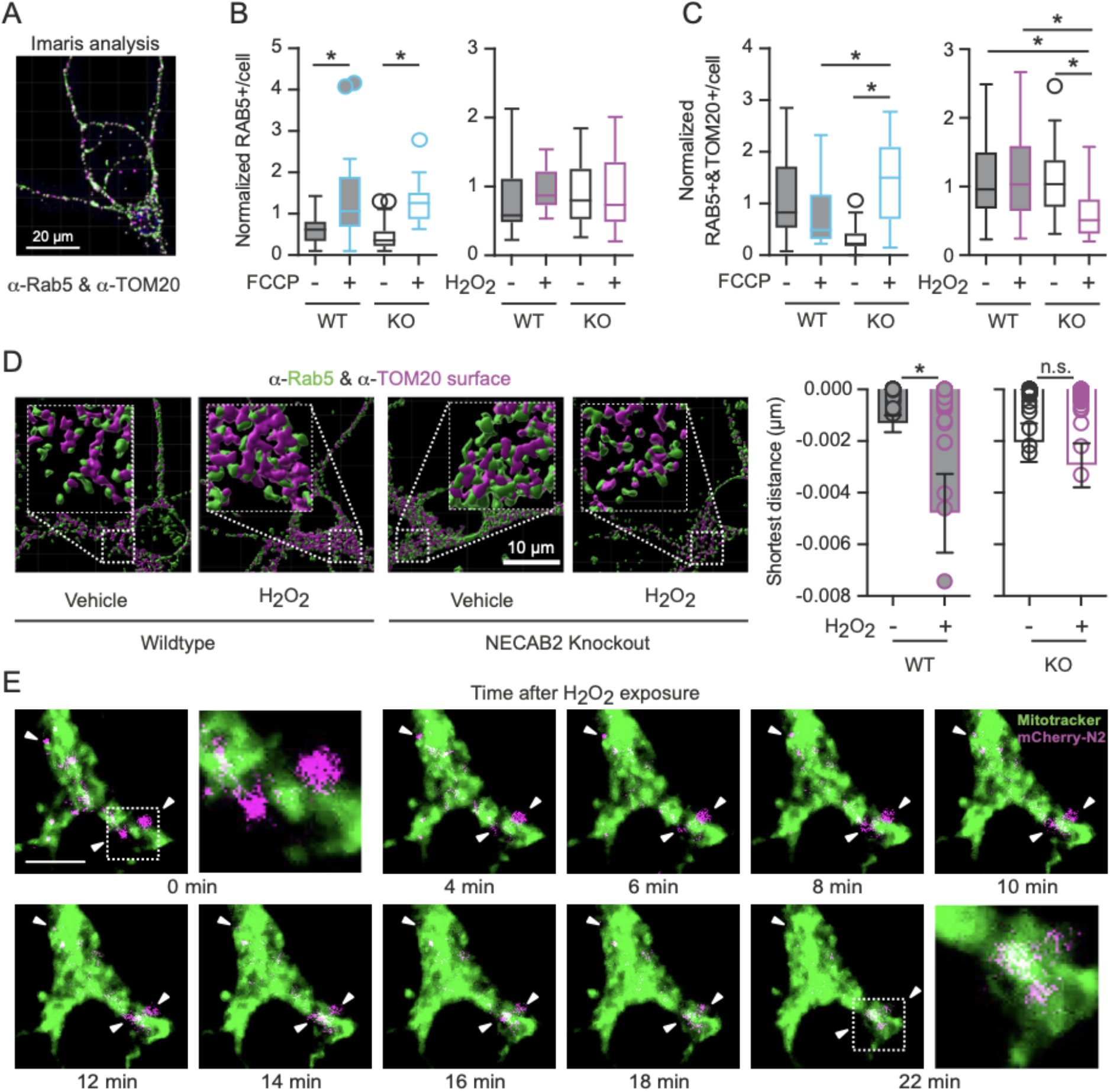
NECAB2 deficiency abrogates endosomal mitochondrial stress response elicited by hydrogen peroxide. (A) Exemplary neuron from primary striatal cultures at DIV 14 stained against Rab5 and TOM20. (B,C) Primary striatal cultures at DIV 12-14 from WT and KO littermates were treated with 10 µM FCCP or 100 µM H_2_O_2_ for 2 h and (B) the number of Rab5 respectively (C) Rab5 and TOM20-positive structures quantified with Imaris. This demonstrated opposing changes in WT and KO neurons. (D) Exemplary neurons from primary striatal cultures at DIV 14 stained against Rab5 and TOM20. Stacked images were used to generate surfaces from the fluorescent signal using the Imaris software. The generated surfaces were used to calculate the distance and the number and distance of contact points between TOM20 and Rab5-positive punctae. 100 µM H_2_O_2_ induces an approximation between the Rab5 and TOM20 punctae only in the WT. Data are presented as individual data points with mean ± SEM or as Tukey plots. (B/C) n=15-37 neurons from 3 mice per genotype, *<0.05 one-way ANOVA. (D) n=48-64 neurons from 3 mice per genotype is indicated, only data points < 0 are shown, *<0.05 Mann Whitney test. *<0.05 Mann Whitney test. (E) Exemplary neuron from primary striatal cultures at DIV 14 transduced with pLV-hSyn-mCherry-NECAB2 five days prior and stained with MitoTrackerGreen. After the addition of 100 µM H_2_O_2_, cells were imaged on a spinning disc confocal microscope at the indicated time points. This demonstrated that mCherry-NECAB2 fuses with mitochondria upon H_2_O_2_ treatment, see white arrowheads and the blown up regions indicated by the dashed lines at the 0 and 22 min timepoints.

We therefore further studied the recruitment of Rab5-positive endosomes to TOM20-positive structures in response to 100 µM H_2_O_2_ by measuring the shortest distance between the surfaces of these structures. Here, negative values denote the presence of TOM20 signal within Rab5-positive structures. WT animals showed the previously described (Hsu et al., 2018) recruitment of Rab5-positive endosomes to mitochondria upon H_2_O_2_ challenge while this was absent in *Necab2* KO striatal neurons (Figure 6D). Interestingly, mCherry-tagged NECAB2 transduced into striatal neurons appears to fuse with mitochondria labeled by mitotracker upon treatment with 100 µM H_2_O_2_ (Figure 6E).

We conclude that NECAB2 is implicated in the endosomal mitochondrial stress response pathway triggered by oxidative signaling and executed by Rab5 and Alsin. NECAB2 fuses with mitochondrial membranes under these conditions and loss of this pathway results in less efficient recycling or repair of mitochondria, synapse loss and striatal dysfunction.

## Discussion

We here describe a novel function for the striatal protein NECAB2. We found that NECAB2 participates in an endosomal mitochondrial stress response mechanism which is important for the correct functioning of the striatum. Without NECAB2, striatal mitochondria are dysfunctional and exhibit increased respiration and superoxide production, resulting in the loss of striatal synapses and behavioral phenotypes related to striatal dysfunction like reduced motivation, increased catalepsy and altered sensory gating.

This mitochondrial stress response mechanism involves the previously reported Alsin-mediated association of Rab5 from early endosomes with mitochondria upon oxidative stress (Hsu et al., 2018). Our study now implicates NECAB2 as a novel player in this process; Alsin and NECAB2 are found in the same macromolecular complex in early endosomes and interact with each other. Without NECAB2 Rab5 does not associate with mitochondria in striatal neurons anymore. Early endosomes are continuously synthesized and exist as a network in the soma of all cells (Zeigerer et al., 2012) and also in dendritic spines (Cooney et al., 2002). Endosomes can therefore serve as a first line of defense when mitochondrial function is compromised or when highly efficient mitochondria are needed. In contrast, autophagosomes, the executors of more canonical mitochondrial quality control, are synthesized *de novo* on demand (Rubinsztein et al., 2012) which does not allow a fast response. The exact function of NECAB2 at the endosome/mitochondria interface and whether it involves its hypothetical enzymatic activity as an atypical heme monooxygenases is however still unclear.

Our data reveal a novel function of NECAB2 that is distinct from its function at the plasma membrane, where NECAB2 boosts basal constitutive signaling of striatal GPCR by direct interaction and increased surface localization (Canela et al., 2009, 2007). We found no alteration in the abundance of these receptors implying that loss of NECAB2 is not compensated by an altered abundance of these receptors. In the striatum, NECAB2 is also not found at the plasma membrane but predominantly intracellular where it strongly colocalizes and co-fractionates with the endosomal protein Rab5. Intriguingly, the interaction of NECAB2 with GPCRs is reversed by increased Ca^2+^ levels (Canela et al., 2009, 2007), making it possible that Ca^2+^ binding results in a conformational change of NECAB2 that might trigger or enhance its localization at intracellular membranes and activation of its function in the endosomal pathway of mitochondrial stress response. Because activation of the NECAB2-interacting metabotropic glutamate receptor mGluR5 results in Ca^2+^ release from the endoplasmic reticulum this would constitute a negative feedback loop. Synaptic stimulation also results in higher cytosolic Ca^2+^ levels that could affect the intracellular localization of NECAB2. Ca^2+^-dependent repair mechanisms involving endosomes have been described at the plasma membrane. Here, assembly of ESCRT proteins at the site of repair is initiated by the Ca^2+^-binding protein ALG-2 (apoptosis-linked gene 2) (Scheffer et al., 2014) leading to pinching out and shedding of wounded membranes (Jimenez et al., 2014). Defects at the plasma membrane clearly increase the local Ca^2+^ concentration, whereas defective mitochondria are rather characterized by excessive production of ROS e.g. the trigger of the Alsin/Rab5-dependent mitochondrial stress response (Hsu et al., 2018). The increased ROS, however, also results in increased Ca^2+^ levels; both alterations amplify each other (reviewed by (Rizzuto et al., 2012)), theoretically allowing a Ca^2+^-dependent recruitment of NECAB2 to mitochondrial membranes. If and how increased Ca^2+^ or ROS levels affect the intracellular localization of NECAB2 needs to be studied further.

NECAB2 is located at striatal synapses where it associates with synaptic mitochondria. *Necab2*-deficient mice have reduced numbers of synapses which correlates with behavioral dysfunction. Because striatal neurons receive massive glutamatergic input from both the cortex and thalamus and dopaminergic input from the midbrain, they have even higher energy demands than other neurons making them more susceptible to mitochondrial dysfunction. Striatal morphology in mice and primates treated with the mitochondrial complex II inhibitor 3-nitropropionic acid closely resembles the features observed here with fewer dendrites and more spines (reviewed by (Brouillet et al., 2005)). Intriguingly, knockout of MIRO1, another EF hand protein that anchors mitochondria at highly active synapses, diminishes the number of dendrites similar to NECAB2 deficiency (López-Doménech et al., 2016). Taken together, mitochondrial maintenance and wellbeing is specifically required at striatal synapses, where NECAB2 is located.

NECAB2 dysfunction might also be involved in human disease. Homozygous Alsin mutation causes familiar ALS, a disease predominantly affecting peripheral and central motoneurons (Weishaupt et al., 2016). In addition to the striatal expression studied here, NECAB2 is also strongly expressed in peripheral α-motoneurons in the ventral horn of the spinal cord (Zhang et al., 2014) making it possible that the NECAB2-mediated mitochondrial stress response is involved in the pathomechanism associated with Alsin mutation. NECAB2 is also strongly expressed in the substantia nigra and degeneration of dopaminergic neurons in the substantia nigra causes Parkinson’s disease (PD). NECAB2 was found to be upregulated in dopaminergic neurons from patients with sporadic PD, monogenic PD associated with mutations in Leucine-rich repeat serine/threonine-protein kinase 2 (LRRK2) (Santiago et al., 2015) and in neurons from patients harboring mutations in the acid β-glucocerebrosidase (GCase) gene (GBA1) (Schöndorf et al., 2014). Mutations in GBA1 constitute the strongest genetic risk factor for PD known to date (Shulman et al., 2011). GCase and LRRK2 both play a role in endomembrane trafficking (Roosen and Cookson, 2016) similar to NECAB2. These results suggest that increased NECAB2 can somehow compensate for the alterations instigated by pathophysiological alterations linked to different PD etiologies. Our data also support the idea that loss of NECAB2 function results in a compensatory upregulation of the recently described ESCRT-dependent pathway of mitochondrial quality control that involves the ubiquitin ligase Parkin (Hammerling et al., 2017) which further links NECAB2 function to Parkinson’s disease. *Necab2*-deficient mice had, however, no motoneuronal or PD-like phenotype suggesting that other cofactors might be involved or that the age when we conducted our behavioral analyses, early adulthood, was too early for any phenotype to become apparent.

In summary, our work has identified a novel player involved in mitochondrial stress response in the striatum. The widespread expression of NECAB2 also suggests a role for this mechanism in other brain regions and in neurodegenerative diseases.

## Materials and Methods

### Ethics

Mice: Animal studies were performed according to the guidelines of the German Animal Welfare Act and Directive 2010/63/EU for the protection of animals used for scientific purposes and was approved by the Rhineland-Palatinate State Authority. Reporting was done in accordance with the Animal Research: Reporting of *In Vivo* Experiments (ARRIVE) guidelines. *Necab2* KO mice of the C57/BL6N background were obtained from KOMP and are named Necab2tm1a(KOMP)Wtsi. The mice were housed in type II long, filter-top cages (Tecniplast, Buguggiate, Italy; SealSafe Plus, polyphenylsulfone, 365mm L x 207mm W x 140mm H Greenline) with enrichment. The mice were maintained under a 12:12-h light/dark cycle (lights on 06:00–18:00) in a temperature- and humidity-controlled animal room (22±2°C, 55±5%). Food (ssniff M-Z Extrudat, ssniff, Soest, Germany) and water were supplied ad libitum.

Human tissue: We used small amounts of anonymized archival surplus post-mortem brain tissue from the Institute of Neuropathology, University Hospital Zurich, Switzerland. The research project did not fall within the scope of the Human Research Act (HRA), thus an authorization of the ethics committee was not required (Cantonal Ethics Committee of Zurich; BASEC-No. Req-2020-00420). [R1]

### Tissue preparation and immunohistochemistry

Brains were perfused with 4% formaldehyde in phosphate buffer saline (PBS), PBS and embedded with paraffin. Coronal sections of 3 µm thickness were cut at the level of the hippocampus using a microtome (Leica Microsystems, Wetzlar, Germany). For immunohistochemistry either paraffin sections that were washed with 0.5% sodium azide (2 × 5 min) or directly cut coronal vibratome sections of 50 µm thickness were used. For the free-floating method the sections were washed several times with PB and blocked with 5% normal goat serum, 0.5% Triton X-100 in the PBS for one hour. Primary antibodies, anti-NECAB2, 1:1000 (Sigma Aldrich, St. Louis, USA), anti-VAMP2, 1:500 (Synaptic Systems, Göttingen, Germany) Alexa-488 tagged cytochrome *c* antibody (BD Biosciences, Franklin Lakes, USA), as well as the secondary antibody goat-anti-rabbit Alexa 568 (1:1000) (Millipore, Burlington, USA) were applied sequentially for 3 h each. After several washing steps with PBS, the sections were stained with 100 nM DAPI for 15 min and then mounted on glass slides with ProLong Gold (Invitrogen, Carlsbad, USA). were acquired using an Ultraview spinning disk confocal microscope (Perkin Elmer, Waltham, USA).

For the NECAB2/VGLUT1/VGLUT2 staining, mice were transcardially perfused with PBS (5 min at 52 rpm) and 4% formaldehyde (PFA) in PBS (5 min at 52 rpm). Brains were then removed and stored at 4°C in PBS with 0.01% sodium azide until cutting. Sections of striatum (bregma coordinates from +1.69 mm to −0.83 mm according to Paxinos and Franklin, 5th edition, 2019), 40 µm thick, were cut with a vibratome (Leica VT1000S). Mouse brain sections were washed in PBS and blocked in 10% normal goat serum (NGS) and 0.1% Triton X-100 in PBS for one hour. The primary antibodies, anti-NECAB2, 1:250, (Sigma Aldrich, St. Louis, USA), anti-VGLUT1, 1:250, (Synaptic Systems, Göttingen, Germany), and Anti-VGLUT2, 1:250, (Synaptic Systems, Göttingen, Germany) were applied in 5% NGS and 0.1% Triton overnight at 4°C. After several washing steps, secondary antibodies (goat-anti-rabbit Alexa 488, (1:1000) and goat-anti-guinea pig Alexa 568, (1:1000) were applied in 5% NGS and 0.1% Triton for one hour at room temperature. DAPI (1:1000) was applied in PBS for an additional 15 min. Sections were washed in PBS, then mounted on a glass slide with Fluoromount G (Thermo Fisher Scientific). Pictures were taken with the confocal multiphoton microscope TCS SP8 (Leica Microsystems, Germany) using 40x and 63x oil immersion objectives.

For the sections of paraffin-embedded mouse brain: Endogenous peroxidase was blocked using 1 ml H_2_O_2_ (30%) in 100 ml methanol/PBS and then incubated for 30 min with 250 µl 5% normal goat serum (NGS) in PBST. Sections were then incubated with the respective primary antibodies overnight at 4°C with polyclonal anti-NECAB2 or polyclonal anti-8-Hydroxydeoxyguanosine. After washing, sections were incubated with Vectastain Kit universal for 30 min, respectively. Afterwards sections were washed, and immunoreactivity was visualized by the avidin-biotin complex method and sections were developed in diaminobenzidine (DAB, Sigma Aldrich, St. Louis, USA).

Human brain sections were dewaxed through graded alcohols and pretreated in LEICA Buffer HIER2 (60 min, 100°C). Sections were automatically stained with Bond autostainer (Leica Microsystems, Wetzlar, Germany) using primary antibody against NECAB2, 1:200 (Sigma Aldrich); for 30 min. Finally, sections were developed using a Bond Polymer Refine Red Detection Kit (Leica Microsystems, Wetzlar, Germany).

### Expansion microscopy of mice brain slices

Brains were processed as indicated above for immunofluorescence with VGLUT1 and NECAB2 antibodies. For secondary antibody incubation we used a goat-anti-guinea pig Alexa 488 (1:1000) and a goat-anti-rabbit Atto 647N (1:1000, #40839, Sigma Aldrich). After antibody incubation, brains were washed trice with PBS for 5 min and free amines were anchored with 0.1mg/mL succinimidyl ester of 6-((Acryloyl)amino)hexanoic acid (Acryloyl-X, SE, Life Technologies) in PBS at 4°C overnight. On the next day, brain slices were washed trice with PBS for 5 min and then transferred into a 12 well plate (CELLSTAR 12 Well plate, Greiner) on ice to initiate gelation. Brains were incubated in 1 mL monomer solution containing 1× PBS, 2 M NaCl, 2.5% (w/w) acrylamide, 0.15% (w/w) N,N’-methylenebisacrylamide, 8.625% (w/w) sodium acrylate, 0.2% (w/w) ammonium persulfate, 0.2% (w/w) tetramethylethylenediamine and 0.01% 4-hydroxy-TEMPO for 30 min at 4°C. For gelation, brains were transferred onto a microscope slide (Menzel Gläser) separated by coverglasses (18×18, 1#, Menzel Gläser) on either side of the tissue, 100 µL of monomer solution was added and a coverglass (18×18, 1#, Menzel Gläser) was placed on top. Gelation was allowed to proceed at room temperature for 2 h. The coverglass was removed with tweezers and the brains were cut out of the gel sheet with a scalpel by leaving 0.5 mm space around. The gel fragments were placed with tweezers in a 50 mL falcon with 1 mL fresh prepared digestion buffer containing 1× TAE buffer, 0.5% Triton X-100, 0.8 M guanidine HCl and 6 units/mL Proteinase K (AM2546, Thermo Fisher). Gel fragments were digested at 4°C overnight. The digestion buffer was replaced by 10 mL Milli-Q water for gel expansion. Milli-Q water was exchanged trice every 30 min until expansion was complete.

Expanded brains were transferred to a staining chamber and embedded with 2% low melting point Agarose in PBS. Confocal immunofluorescence was performed with the TCS SP8 (Leica Microsystems, Germany) using 40x and 63x oil immersion objectives.

### Immunocytochemistry

Striatal neurons were plated on coverslips coated either with a mixture of 0.5 mg/ml PDL solution (Sigma Aldrich, St. Louis, USA), 1 mg/ml collagen solution (Thermo Fisher Scientific, Waltham, USA) and 17 mM acetic acid solution or with PLL solution (0.1 mg/ml). DIV 12 MSNs were fixed with 4% Roti®-Histofix (20 min at RT) and permeabilized with 0.25% Triton X-100 (Thermo Fisher Scientific, Waltham, USA) in PBS for 10 min at RT. The coverslips were blocked with Roti®-ImmunoBlock 1X (Carl Roth, Karlsruhe, Germany) for 30 min to 1 hr, followed by incubation with primary antibodies anti-NECAB2, 1:500 (Sigma Aldrich, St. Louis, USA), anti-Rab5, 1:200 (Santa Cruz, Dallas, USA), anti-Tom20, 1:400 (Sigma Aldrich, St. Louis, USA), anti-HSPA9, 1:50 (NeuroMab, Davies, USA), anti-CD63, 1:50 (Santa Cruz, Dallas, USA), anti-LC3B, 1:100 (NanoTools, Teningen, Germany), anti-PDI, 1:200 (Cell Signalling, Danvers, USA) at 4°C with gentle shaking overnight. The neurons or the transfected cells were washed and reprobed with fluorescent conjugated secondary antibodies (anti-Rabbit AlexaFluor-568 1:1000 (Life Technologies, Carlsbad, USA) and anti-Mouse AlexaFluor-488 1:000 (Life Technologies, Carlsbad, USA) or anti-Mouse AbberiorSTAR440 1:200 (Abberior, Göttingen, Germany) and anti-Rabbit Chromeo505 1:200 (Active Motif, Carlsbad, USA)) in 1X Roti®-ImmunoBlock in dark condition at room temperature for 1 hr. Coverslips were stained with 300 nM DAPI for 5 min at RT and mounted onto microscope slides with Dako Fluorescence Mounting Medium. Fluorescence pictures were taken either with a confocal microscope (TCS SP5 or TCS SP8 – Leica Microsystems, Wetzlar, Germany) or with a Zeiss Axiovert 200 M (Carl Zeiss, Oberkochen, Germany) with deconvolution software using a 100x/1.4 oil immersion objective. STED images were acquired with a TCS STED CW super-resolution microscope using a 100x/1.4 oil immersion objective. Colocalization studies were performed using the JACoP plugin from ImageJ (Bolte and Cordelières, 2006) and/or using Imaris (BitPlane, Belfast, UK), where 3D images were generated from z-stacks. The colocalization degree of both channels were determined followed by quantification of dots of colocalizing signal, after applying a Background Subtraction (1.0) and Gaussian (0.1) filters. Alternatively, surfaces were generated from the fluorescent signal (threshold diameter size for mitochondria: 0.3 μm; endosomes: 0.4 μm; NECAB2 vesicles: 0.6 μm) on Imaris and the shortest distance between surfaces and the number of surfaces in contact were quantified.

### Live cell imaging

WT primary striatal neurons were plated on pre-coated 8-well Ibidi slides (Ibidi, Gräfelfing, Germany). On DIV 9, neurons were transduced with a pLV-hSyn-mCherry-NECAB2 lentivirus. After 5 days, the neurons were stained with MitoTracker Green (Thermo Fisher Scientific, Waltham, USA) and imaged with a Visitron Spinning Disc microscope (Visitron, Puchheim, Germany) in a humidified atmosphere at 37°C and 5% CO_2_. After the addition of 100 µM H_2_O_2_, the neurons were imaged every 2 min for 30 min.

### Immunoblotting

Denatured total cellular protein samples (95°C, 5 min) from cells or brain tissue were separated on SDS polyacrylamide gels (4–15% Mini-PROTEAN® TGX Stain-Free™ gels, Bio-Rad, Hercules, USA) and transferred onto a membrane (nitrocellulose or PVDF) using the iBlot Dry Blotting System (Invitrogen, Carlsbad, USA) or Trans-Blot® Turbo™ Transfer System (Bio-Rad, Hercules, USA). Membranes were blocked with 3 or 5% (w/v) milk powder in PBS, PBS-T (1x PBS, 0.05% (v/v) Tween 20) or TBS-T (1x TBS, 0.05% (v/v) Tween 20) for 1 h at room temperature (RT). The membranes were incubated with the primary antibodies anti-NECAB2 1:1000 (Sigma Aldrich, St. Louis, USA), anti-VDAC 1:1000 (Cell Signaling, Danvers, USA), anti-COXIV 1:1000 (Cell Signaling, Danvers, USA), anti-b-tubulin 3, 1:2000 (R&D Systems, Minneapolis, USA),, anti-PSD95, 1:1000 (Synaptic Systems, Göttingen, Germany), anti-b-actin mAB, 1:4000 (Millipore, Burlington, USA), anti-Rab5, 1:1000 (Santa Cruz, Dallas, USA) respectively, overnight at 4°C. For visualization, membranes were incubated with infrared fluorescence IRDye 680 and 800-conjugated anti-mouse/anti-rabbit IgG secondary antibodies, 1:30000 (LI-COR Biosciences, Lincoln, USA) for 1 h at RT and detected with the Odyssey Infrared or Odyssey Sa Infrared Imaging System (LI-COR Biosciences, Lincoln, USA). The software ImageJ or Image Studio™ Lite Software (https://www.licor.com/bio/image-studio-lite/) was used to analyze the expression of proteins in relation to the control.

### Autoradiography

Animals were decapitated and their brains were rapidly removed, blotted free of excess blood, immediately frozen in 2-methylbutane (−50°C) and stored until sectioning. Coronal sections (20 µm thickness) were produced in a microtome (Leica Microsystems, Wetzlar, Germany), thaw-mounted onto silane-coated slides and stored at −80°C until further use. All binding assays were preceded with a preincubation in the respective buffers for 15-30 min to remove endogenous transmitters. Adenosine A2A, dopamine D1 and D2 and metabotropic glutamate mGluR5 receptors as well as dopamine transporters (DAT) were labeled accordingly with [^3^H]ZM241385 (0.42 nM), [^3^H]SCH23390 (1.61 nM), [^3^H]raclopride (0.59 nM), [^3^H]ABP688 (2.23 nM) and [^3^H]mazindol (4.07 nM) in 50 mM Tris-HCl (pH 7.4; 90 min at 22°C) containing in the pre- and main incubation EDTA (1 mM) buffer. Additionally, the main incubation adenosine deaminase (2 U/mL) in 50 mM Tris-HCl (pH 7.4; 90 min at 22°C) contains NaCl (120 mM), KCl (5 mM), CaCl_2_ (2 mM) and MgCl_2_ (1 mM). Furthermore, the main incubation mianserin (1 µM) in 50 mM Tris-HCl (pH 7.4; 45 min at 22°C) contain NaCl (150 mM) and ascorbic acid (0.1%) in 30 mM HEPES (pH 7.4; 65 min at 22°C) contain NaCl (40 mM), KCl (5 mM), CaCl_2_ (2.5 mM) and MgCl_2_ (1.2 mM) respectively or in the preincubation buffer 50 mM Tris-HCL (pH 7.9, 60 min at 4°C) contain 120 mM NaCl and 5 mM KCL and in the main incubation buffer 300 mM NaCl, 5 mM KCL with 300 nM desipramine using the displacers theophylline (5 mM), SKF 83566 (1 µM), butaclamol (1 µM), MPEP (10 µM) and nomifensine (100 µM), respectively. Brain tissue slices were washed two to six times in ice-cold buffer and rapidly rinsed in ice-cold distilled water, placed under a stream of cold air and exposed to phosphor-imaging plates (BAS-TR2025, Raytest, Straubenhardt, Germany) for three days together with tritium standards. Autoradiograms were scanned with a high-performance plate reader (BAS-5000, Raytest, Straubenhardt, Germany) and subsequently processed in a blinded manner using image-analysis software (AIDA 4.13, Raytest, Straubenhardt, Germany).

### Primary neuronal culture

Primary striatal neurons were cultured from embryonic day 17 (E17). Embryos were obtained from pregnant NECAB2 heterozygous females. *Necab2* KO and WT brains were removed from the skull and placed in cold Hank’s buffer without Ca^2+^ and Mg^2+^ (HBSS -/-). The cerebellum and brainstem were separated. Under a dissecting microscope striatal tissue was dissected and pooled separately after removal of the meninges. Dissected tissues of each embryo were kept separately, and the tails were collected for post genotyping. Tissues were washed once with HBSS -/- buffer, replaced with fresh 5 ml of HBSS -/- and digested for 15 min at 37ºC with trypsin/EDTA (10X, Sigma Aldrich, St. Louis, USA). DNaseI (Sigma Aldrich, St. Louis, USA) (50 g/ml) was also added at the same time of trypsinization. Tissues were then washed twice with plating media containing serum [Neurobasal medium (Thermo Fisher Scientific, Waltham, USA) supplemented with 2% B27 (Thermo Fisher Scientific, Waltham, USA), 1% Glutamax (Thermo Fisher Scientific, Waltham, USA), 1 μg/ml penicillin/streptomycin (Sigma Aldrich, St. Louis, USA) and 5% FCS (Thermo Fisher Scientific, Waltham, USA)] to quench the trypsin and dissociated into a single-cell suspension by gentle repeated passages through a sterile glass pipette.

For biochemical experiments, striatal neurons were plated at a density of 50K per well onto 24 well plates or 20K per well onto 8 well chamber slides coated with poly-L-lysine (Sigma Aldrich, St. Louis, USA) (100 µg/ml). Cells were maintained in plating media with serum for 3-4 h and later replaced with Neurobasal media without serum [supplemented with 2% B27, 1% Glutamax and 1 μg/ml penicillin /streptomycin]. Neurons were incubated at 37°C with 5% CO2. A solution of 10 mM 5-fluoro-2’-deoxyuridine and uridine (anti-mitotic agent) (Sigma Aldrich, St. Louis, USA) was added to the cultures at days in vitro (DIV) 3-6 to prevent glial overgrowth.

### Subcellular fractionations

Endosomal fractions: Brains were removed from the *Necab2* KO and WT mice. The striatum was isolated and washed once with 5 ml PBS. After removing the supernatant, endosomal fractions were prepared as described in (Aniento and Gruenberg, 2008; Scheffer et al., 2013). The cell pellet was gently resuspended in 1.8 ml of homogenization buffer (HB: 250 mM sucrose, 3 mM imidazole, pH 7.4, 10 mM Tris–HCl, pH 7.4) using a wide pore tip (2 mm diameter). Afterwards, the homogenate was transferred to a 2 ml reaction tube and centrifuged for 10 min at 1500 x g at 4°C. The supernatant was removed, and the pellet was resuspended in 0.5 ml of HB using a blue tip (1 ml). Immediately, the homogenate was quickly drawn ten times through a 22-G needle attached to a 1 ml syringe. The homogenate was centrifuged for 10 min at 1500 x g at 4°C. The post-nuclear supernatant (PNS) was taken for the sucrose gradient and diluted with a 62% sucrose solution (2.351 M sucrose, 3 mM imidazole, pH 7.4, 1 mM EDTA) to obtain a final concentration of 40.6%. The PNS sample was loaded to the bottom of a SW60 centrifuge tube and was overlaid with a 35% sucrose solution (1.177 M sucrose, 3 mM imidazole, pH 7.4). The next layer was a 25% sucrose solution (0.806 M sucrose, 3 mM imidazole, pH 7.4) and the homogenization buffer was used to fill up the centrifuge tube. The sucrose concentration was adjusted by monitoring the refractive index of each solution (1,3723 for 25%; 1,3904 for 35%; 1,4463 for 62%; 1,4001 for 40,6%). After centrifuging in a SW60 rotor for 90 min at 14000 x g, the gradient was fractionated in 300 µl steps. Early endosomes were enriched at the 25%/35% interface and late endosomes were located at the 25%/HB interface. The fractions were analyzed by immunoblotting (see above) and isolated endosomes were used for subsequent proteomics analyses.

Synaptic and non-synaptic mitochondria: All the steps were carried out at 4°C with extreme caution. Mouse brains were quickly removed and homogenized in a 4 ml Mito I buffer (0.32 M sucrose, 10 mM Tris-HCl pH 7.4, 1 mM EDTA, protease and phosphatase inhibitors) using a loosely fitting glass potter. Homogenized brains were further disrupted for 10 min using a cell disrupting bomb (Parr Instrument Company, Frankfurt am Main, Germany) by applying pressure of 60 bar of nitrogen gas, followed by the protocol as previously described in (Wieckowski et al., 2009) for isolation of mitochondria from animal tissue. Concisely, the homogenate was centrifuged with increasing speed and mitochondria were finally purified using a Percoll gradient method. The nuclear, cytosolic, crude and pure mitochondrial fractions were used for further analyses. Synaptic and nonsynaptic mitochondria were prepared adapting previously published protocols. Striata from 5 mice were pooled and homogenized with a Dounce homogenizer in a 5x volume of ice-cold isolation buffer containing 215 mM mannitol, 75 mM sucrose, 20 mM HEPES and 1mM EGTA, pH: 7.2. The resultant homogenate was layered on a discontinuous gradient of 15%, 23%, and 40% (vol/vol) Percoll and centrifuged at 34,000 x g for 10 min. After centrifugation, band 2 (the interface between 15% and 23% containing synaptosomes) and band 3 (the interface between 23% and 40% containing nonsynaptic mitochondria) were removed from the density gradient. The fractions were washed by centrifugation at 16,500 x g for 15 min. The pellets were collected and resuspended in isolation buffer and the synaptosomal fraction was disrupted for 10 min using a cell disrupting bomb (Parr Instrument Company, Frankfurt am Main, Germany) by applying pressure of 1000 psi for 10 min. A second Percoll density gradient was prepared and the samples were layered for ultracentrifugation as described above. Band 3 was obtained and resuspended in isolation buffer and the pellets were washed three times in isolation buffer without EGTA. The final synaptic and nonsynaptic mitochondrial pellets were resuspended in isolation buffer and stored on ice.

Synaptosomes: After homogenizing the brain tissue with 10 ml/1 g 0.32 M sucrose, 5 mM HEPES (pH 7.4), and protease inhibitor, the homogenate was centrifuged for 10 min, 1000 x g at 4°C and the supernatant (S1) was collected. The remaining pellet (P1) was homogenized as in the first step and then it was again centrifuged and the supernatant (S1’) was collected. The supernatants S1 and S1` were combined and centrifuged at high speed for 15 min, 12,000 x g at 4°C. The supernatant (S2) was removed. The pellet included the synaptosome crude extract (P2). After washing, the pellet was dissolved and centrifuged for 20 min, 12,000 x g at 4°C. The supernatant (S2`) was removed. The pellet included synaptosome crude extract (P2). After resuspending the pellet in 1.5 ml/1 g 0.32 M Sucrose, 5 mM HEPES (pH 8.1), and protease inhibitor, the pellet was loaded on a sucrose step gradient (SW40Ti: 10 ml tubes, 3 ml per gradient). Afterwards the gradient was centrifuged for 2 h, 26,000 rpm at 4°C in a Beckman Coulter Ultracentrifuge and the visible synaptosome layer was collected from the layer between 1 and 1.2 M sucrose. The synaptosome fractions were used for Western blot and electron microscopic analyses.

### Transmission electron microscopy and image quantification

Synaptosomes were fixed with 2% glutaraldehyde in 0.1 M sodium phosphate buffer, pH 7.4. After washing, the samples were post-fixed with 2% osmium tetroxide/1.5% potassium ferrocyanide for 1 h on ice, washed, contrasted en bloc with uranyl acetate, dehydrated with an ascending series of ethanol and embedded in glycid ether 100-based resin. Ultrathin sections were cut with a Reichert Ultracut S ultramicrotome (Leica Microsystems, Wetzlar, Germany). After contrasting, they were viewed with a ZEISS EM 10 CR electron microscope at an acceleration voltage of 60 kV. The length, width and electron density of the mitochondria were analyzed with Fiji/ImageJ.

TEM of brain striatum was performed as described previously (Anand et al., 2016). Mice were fixated by performing whole body cardiac perfusion using freshly prepared 3.7% formaldehyde and 0.5% glutaraldehyde buffered in PBS. Perfused brain tissue was washed twice with 0.1 M sodium cacodylate buffer (pH 7.2). Small striatal tissue pieces of about 2 mm^3^ size were prepared. Striatal tissue pieces were stained for 1 h with 1% OsO_4_ in 0.1 M sodium cacodylate buffer (pH 7.2). Samples were washed three times for 10 min with aqua bidest prior to staining with 1% uranyl acetate in aqua bidest at 4°C overnight. Stained pieces were washed three times for 5 min with aqua bidest before subjecting them to a series of ethanol dehydration incubations: 1 × 20 min in 30% ethanol, 1 × 20 min in 50% ethanol, 1 × 30 min in 70% ethanol, 1 × 30 min in 90% ethanol, 1 × 30 min in 96% ethanol, 1 × 30 min in 100% ethanol. This was followed by resin infiltration incubations: 1 × 20 min in 50% resin (AGR1280; Agar Scientific, Stansted, UK) in 100% ethanol, 3 × 20 min in 100% resin. Finally, specimens were embedded in 100% resin in gelatin capsules and incubated for polymerization at 65°C for 16 h. A transmission electron microscope (Hitachi, H600) was used for image recording of ultrathin sections. Recordings were made at 75 V equipped with Bioscan model 792 (Gatan) camera.

Electron microscopic images of striatal mitochondria were analyzed using ImageJ. Each single image had a size of 1 µm^2^. Length and width of each mitochondrion was measured by defining the spatial scale and also by measuring “Distance in Pixels”. Each mitochondrion was identified and counted manually as regions of interest (ROI) and the average signal intensity (mean grey value) within the ROI was measured. The mean grey value of each selected ROI referred to the levels of density of each mitochondrion. Synapses were counted manually (blind and unbiased) and divided by the total area to obtain the number of synapses per µm^2^.

### High resolution respirometry

Mice were sacrificed by cervical dislocation immediately before analysis. Brains were micro-dissected on ice and dissected striata were directly transferred into ice-cold mitochondrial respiration medium MiR05 (Gnaiger et al., 2000). The striatum was then homogenized in a pre-cooled 1.5 ml tube with a motorized pestle. Resulting homogenates containing 10 mg tissue w/w were suspended in 100 ml of MiR05 and later 20 ml (2 mg) tissue suspension was added to each chamber of the Oroboros Instrument (Oroboros Instruments, Innsbruck, Austria) containing 2 ml of MiR05 for OXPHOS-analysis (Holmström et al., 2012). Oxygen concentration (μM) as well as oxygen flux per tissue mass (pmol O2. s^−1^·mg^−1^) were recorded in real-time using DatLab software (Oroboros Instruments, Innsbruck, Austria). A multi substrate protocol was used to sequentially explore the various components of mitochondrial respiratory capacity. The striatum homogenate was suspended in MiR05, added to the Oxygraph-2k glass chambers and the O_2_ flux was allowed to stabilize. A substrate, uncoupler, inhibitor titration (SUIT) protocol was applied to assess qualitative and quantitative mitochondrial changes in NECAB2 KO mice and WT controls. After stabilization, LEAK respiration was evaluated by adding the complex I (CI) substrates malate (2 mM), pyruvate (10 mM) and glutamate (20 mM). Maximum oxidative phosphorylation (OXPHOS) capacity with CI substrates was attained by the addition of ADP+MgCl_2_ (5 mM) (CI OXPHOS). For evaluation of maximum OXPHOS capacity of the convergent input from CI and complex II (CII) at saturating ADP concentration, the CII substrate succinate (10 mM) was added (CI+CII OXPHOS). We then uncoupled respiration to examine the maximal capacity of the electron transport system (ETS or CI+II ETS) using the protonophore, carbonyl cyanide 4 (trifluoromethoxy) phenylhydrazone (FCCP; successive titrations of 0.2 μM until maximal respiration rates were reached). We then examined consumption in the uncoupled state solely due to the activity of complex II by inhibiting complex I with the addition of rotenone (0.1 μM; ETS CII or CII ETS). Finally, electron transport through complex III was inhibited by adding antimycin A (2 μM) to obtain the level of residual oxygen consumption (ROX) due to oxidation side reactions outside of mitochondrial respiration. The O_2_ flux obtained in each step of the protocol was normalized by the wet weight of the tissue sample used for the analysis and the ROX was subtracted from the fluxes in each run to correct for non-mitochondrial respiration (Hollis et al., 2015).

### Superoxide quantification

The brains of WT and *Necab2* KO mice were removed and one part of the striatum was isolated. The striatum tissue was resuspended in 500 µl PBS. After centrifugation, the supernatant was discarded, and the tissue resuspended and incubated in 20 µM myxothiazol solution in PBS (100 µl) for 20 min at RT. After another centrifugation step, the supernatant was again discarded, and the tissue washed with 500 µl PBS. The tissues were incubated with 50 µM dihydroethidium (200 µl) for 30 min at RT in the dark. After centrifugation, the supernatants were collected for control measurements while the tissue pellet was resuspended in 200 µl PBS and stored on ice. The samples were diluted 1:1 with 100% acetonitrile and then centrifuged for 20 min at 20,050 x g. 200 µl of the supernatant were transferred to a high-performance liquid chromatography (HPLC). The method was described in (Schuhmacher et al., 2011). DHE signal was detected by its absorption at 355 nm. The DHE signal was normalized to the wet weight of the striatum tissue.

### Enzymatic measurement of total cellular GSH content

Striata were isolated from 3 or 12-month-old *Necab2* KO and WT mice. Striata were homogenized in 200 µl of 2 mM EDTA in cold PBS each in separated 1.5 ml tubes. Total cellular GSH content was quantified enzymatically as previously described (Lewerenz et al., 2006). The tissue homogenate was carefully washed twice with phenol red-free Neurobasal medium and incubated with 20 µM MCB for 5 min at 37°C protected from light. MCB fluorescence was measured by a SpectraMax I3 microplate reader. Excitation wavelength was 393 nm and emission wavelength 485 nm.

### Behavioral tests

Fifteen male WT and *Necab2* KO mice (littermates) at 3-4 months of age underwent a battery of behavioral tests. All tests were performed in sound-attenuated rooms, between 9 am and 5 pm. Mice were grouped with access to food and water and a 12 h - 12 h light-dark cycle (light phase onset at 7.00 h). All experiments were performed with permission of the local authorities in accordance with the German Animal Protection Law. All experiments and initial quantitative analyses were performed blinded, i.e. the experimenter had no knowledge of the genotype of the mice tested and analyzed.

Progressive ratio test: Progressive ratio reinforcement task to evaluate motivation: Progressive ratio (increasing number of responses are required to obtain consecutive deliveries of the reward) reinforcement task provides reliable and unambiguous measure of incentive motivation. We conducted this task in the touch-screen-based operant conditioning system (Med Associates, St. Albans, VT, USA). The operant chamber (dimensions: 18.1 cm × 17.6 cm × 18.6 cm), enclosed in a sound and light attenuating box, contained a pellet dispenser, which delivered a single 14 mg dustless precision pellet (Test Diet, St. Louis, USA) into a food tray (magazine). A touch sensitive TFT monitor (touchscreen; 18.4 cm × 13.6 cm) was located on the opposite side of the chamber. The touchscreen was covered with a metal partition containing a response hole (2 cm in diameter), allowing presentation of simple visual stimuli (light). Touch-screen-based progressive ratio reinforcement task was controlled by K-Limbic software (Conclusive Marketing Ltd, Herts, UK).

Touch-screen experimental procedures: Initially, a food deprivation regimen was introduced, and the mice were gradually reduced to 85% of their free-feeding body weights. Animals were kept on food deprivation regimen throughout the duration of the experiment. Mice were tested on 7 consecutive days per week. Every mouse underwent the following sequence of test phases: (1) reward habituation and chamber acclimation, (2) autoshaping, (3) pre-conditioning 1–3 and (4) progressive ratio reinforcement task. Daily sessions were terminated after 30 min or as soon as the mouse had accomplished the learning criterion of the respective test phase.

Reward habituation and chamber acclimation: First, mice were habituated overnight with reward pellets in their home cages. Next day an experimental chamber acclimation began: Pellets were freely available in the magazine with an inter-trial interval of 15 s. A new trial was initiated after collection of the pellet leading to a disruption of an infrared light beam. This phase was conducted daily for 30 min until the mouse reached the criterion: No pellets left in the magazine after finishing the session. Then the mouse was moved to the next phase.

Fixed ratio 1 (FR1): The session started with a reward pellet in a magazine. After the mouse had collected the pellet, a light stimulus was presented in the stimulus hole on the touchscreen. Stimulus was presented until touch-response of the mouse. Then, the stimulus light went OFF and light in the magazine went ON, signalizing delivery of the reward pellet. A next trial was initiated after collection of the pellet (inter-trial interval 15 s). This phase was conducted daily for 30 min until the mouse reached the criterion: Complete at least 30 trials, and then the mouse was moved to the next phase.

Fixed ratio 2 (FR2): This phase was performed exactly as FR1, with one exception: The mouse has to make 2 touch-responses in order to be rewarded. After reaching the criterion, the mouse was moved to the next phase.

Fixed ratio 4 (FR4): This phase was performed exactly as FR1, with one exception: The mouse has to make 4 touch-responses in order to be rewarded. After reaching the criterion, the mouse was moved to the final phase.

Progressive ratio (PR) (Kheramin et al., 2005). This phase was performed similarly as FR1. Progressive ratio means that a mouse has to progressively increase the number of responses (touches of the screen) in order to earn a reward pellet. The PR schedule (i.e. the number of responses mice have to do in order to get next reward) was based on the following exponential progression: 1, 2, 4, 6, 9, 12, 15, 20, 25, 32, 40, 50, …, derived from the formula [(5×e0.2n)−5], rounded to the nearest integer, where “n” is the position in the sequence of ratios. The duration of this phase was 90 min. A number of completed (i.e. rewarded) trials was recorded for each mouse.

Catalepsy test: The bar test was adapted from Meschler *et al*. (Meschler et al., 2000). A metal bar (4mm in diameter) was horizontally positioned 6 cm above the desk surface. The animal’s front paws were placed on the bar and the hind paws were rested on the desk. Time was recorded until the mouse placed both front paws onto the desk.

Prepulse inhibition test of sensory-motor gating (Koch, 1999): The test was performed as described earlier (Radyushkin et al., 2009). Briefly, mice were placed in small metal cages (90 × 40 × 40 mm) to restrict major movements. The cages were equipped with a movable platform floor attached to a sensor that records vertical movements of the floor. Cages were placed in four sound-attenuating isolation cabinets (Med Associates, St. Albans, USA).

Startle reflexes were evoked by acoustic stimuli delivered from a loudspeaker suspended above the cage and connected to an acoustic generator. The startle reaction to an acoustic stimulus, which evokes a movement of the platform and a transient force resulting from this movement of the platform, was recorded with a computer during a recording window of 380 milliseconds (beginning with the onset of pre-pulse) and stored for further evaluation. An experimental session consisted of a 2-min habituation to 65 dB background white noise (continuous throughout the session), followed by a baseline recording for 1 min. After baseline recording, 8 pulse-alone trials using startle stimuli of 120 dB intensity and 40 milliseconds duration were applied in order to decrease the influence of within-session habituation. These data were not included in the analysis of the prepulse inhibition. For tests of prepulse inhibition, the 120 dB/40 milliseconds startle pulse was applied either alone or preceded by a pre-pulse stimulus of 70, 75- or 80-dB intensity and 20 milliseconds duration. An interval of 100 milliseconds with background white noise was employed between each pre-pulse and pulse stimulus. 10 trials of each kind (in total 40 trials) were applied in a pseudorandom order with variable inter-trial intervals ranging from 8 to 22 seconds. Amplitudes of the startle response were averaged for each individual animal, separately for each type of trials (i.e. stimulus alone or stimulus preceded by a pre-pulse f 3 intensity). Prepulse inhibition was calculated as the percentage of the startle response using the following formula: PPI (%) = 100 – [(startle amplitude after pre-pulse and pulse / startle amplitude after pulse only) × 100].

Open field test as described (Jamain et al., 2008): Instinctive movement in the open field was tested in a gray Perspex arena (120 cm in diameter, 25 cm high; illumination 120 lux). Mice were placed in the center and allowed to explore the open field for 7 min. The behavior was recorded by a PC-linked overhead video camera. “Ethovision XT” (Noldus Information Technology, Wageningen, Netherlands), a video-tracking software, was used to calculate velocity, distance traveled, and time spent in central, intermediate or peripheral zones of the open field.

Elevated plus maze as described (Jamain et al., 2008): Individual mice were placed in the central platform facing an open arm of the plus-maze (made of grey Perspex with a central 5 × 5 cm central platform, two open arms, 30 × 5 cm, 2 enclosed arms, 30 × 5 × 15 cm, illumination 120 lux). Behavior was recorded by an overhead video camera and a computer equipped with “Ethovision XT” (Noldus Information Technology, Wageningen, The Netherlands) software to calculate the time each animal spent in open or closed arms. The relevant amount of time spent in open arms was used to evaluate open arm aversion, which is associated with fear.

### Golgi-Cox staining and analysis

For Golgi-Cox analysis the PBS-perfused brains of WT (n=7) and *Necab2* KO (n=5) mice were processed as described (Frauenknecht et al., 2015). A LEICA DM6000B microscope and a computer-based system (Neurolucida; MicroBrightField Bioscience, Williston, USA) were used to generate three-dimensional neuron tracings. Visualization and analysis of neuron morphology were performed using NeuroExplorer (MicroBrightField Bioscience, Williston, USA). Neurons were reconstructed by one investigator blinded to group identity using a camera lucida and a 40x objective. Dendritic parameters as well as 3D Sholl analysis to assess dendritic complexity were calculated for each neuron and for each animal as described (Frauenknecht et al., 2015). The results were analyzed for apical and basal dendrites separately and were presented as means ± SEM. The density of dendritic spines of a distinct branch order (MSN: 2nd, 3rd and 4th order dendrites) was measured for each animal using a 100x oil-immersion objective (Leica Microsystems, Wetzlar, Germany). Dendritic spines visible along both sides of segments were counted on dendrites of distinct orders with a minimum length of 10 µm and were expressed as mean number of spines per µm.

### *In utero* electroporation and spine quantification

*In utero* electroporation experiments were carried out essentially as described (Baumgart and Grebe, 2015). Injection was made into the striatum. After birth, the electroporated mice were decapitated and perfused with 4% PFA at the age of P21. Staining on free-floating vibratome slices (40 µm) was done with anti-GFP (Abcam, Cambridge, UK). Pictures were taken with a 63x oil immersion objective on the confocal microscope TCS SP5 from Leica (Leica Microsystems, Wetzlar, Germany). Analysis and 3D construction were done with ImageJ and Imaris.

### Quantification and statistical analysis

The Shapiro-Wilk test was used to determine normality of distribution of the data. Data normally distributed were analyzed with the one-or two-way ANOVA followed by the Tukey’s test or with the independent t test. Data not-normally distributed were analyzed with the Kruskal-Wallis test followed by the Dunn’s test or with the Mann-Whitney test. A *P* value lower than or equal to 0.05 was considered statistically significant. Values are reported as mean ± SEM. Additional analyses were performed using the general statistics module of Analyse-it™ for Microsoft Excel (Analyse-it Software Ltd., Leeds, UK) or by using GraphPad Prism (GraphPad, San Diego, USA).

## Acknowledgment

This work was supported by the Deutsche Forschungsgemeinschaft (ME1922/14-1) to AM and the German Ministry of Education and Research (BMBF; GeNeRARe 01GM1519A) to MJS. The IMB Core Facility Microscopy and the Core facility for electron microscopy (CFEM, Heinrich Heine University Düsseldorf, Medical faculty) are gratefully acknowledged for their support. We are grateful to Hilmar Bading, Department of Neurobiology and Interdisciplinary Center for Neurosciences, Heidelberg University, in whose laboratory the electron microscopy work of the synaptosomes was carried out. The article contains results of the MD theses of Elena Morpurgo and Lena Sessinghaus. We thank Elizabeth Jane Rushing for excellent proofreading.

## Author contributions

Performed experiments: PND, DB, TS, CW, VW, EM, LR, LS, PL, KR, MKES, LF, JB, PS, GH, LN, VV, AH, AO, NN, RA. Analyzed data and wrote materials and methods: PND, DB, TS, CS, KR, LF, CJ, JB, AD, AB, ASR, SR, JP, MS, KBMF, AM. Wrote the manuscript: MS, AM

## Data availability

All data needed to evaluate the conclusions in the paper are present in the paper and/or the Supplementary Materials.

## Supplemental figures

**Supplemental Figure 1:**
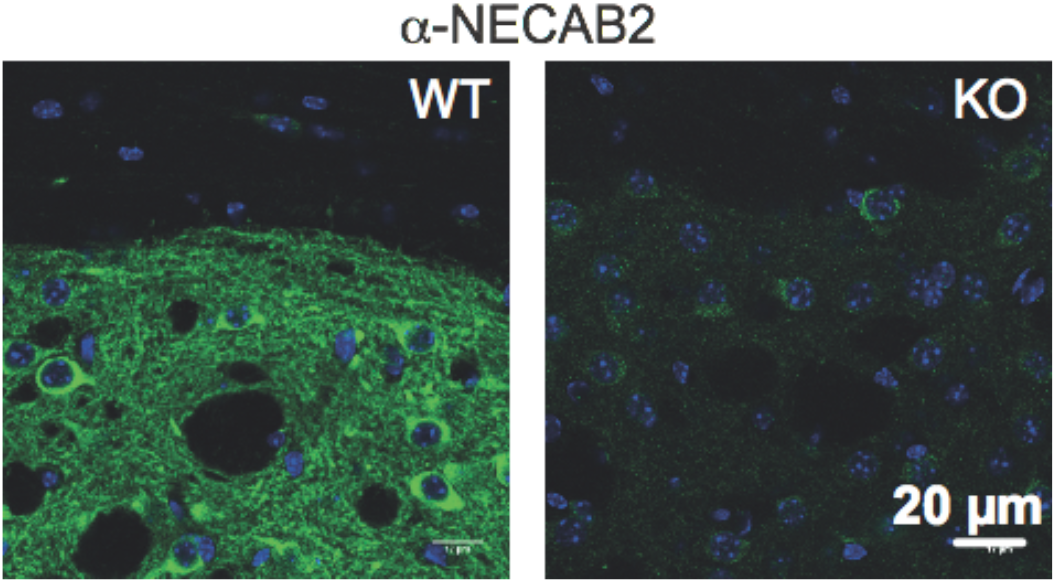
Specificity of the NECAB2 antibody shown by immunohistochemistry of wildtype (WT) and knockout (KO) striatal brain sections. Size is indicated.

**Supplemental Figure 2:**
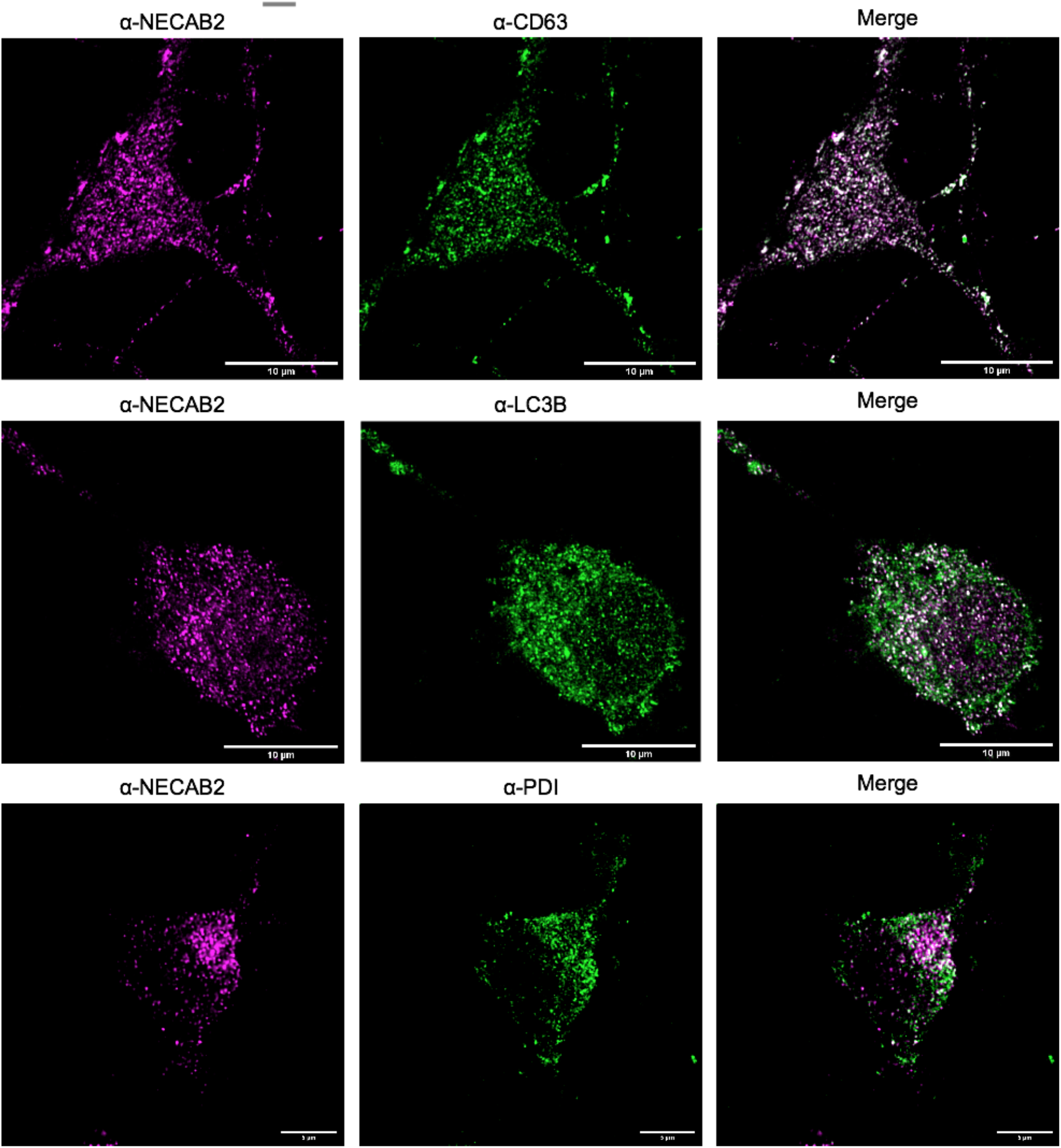
NECAB2 expression partially overlaps with CD63, a marker for multivesicular bodies, LC3, a marker for lysosomes, and PDI as a marker for the endoplasmic reticulum. Size marker 10 µm.

**Supplemental Figure 3:**
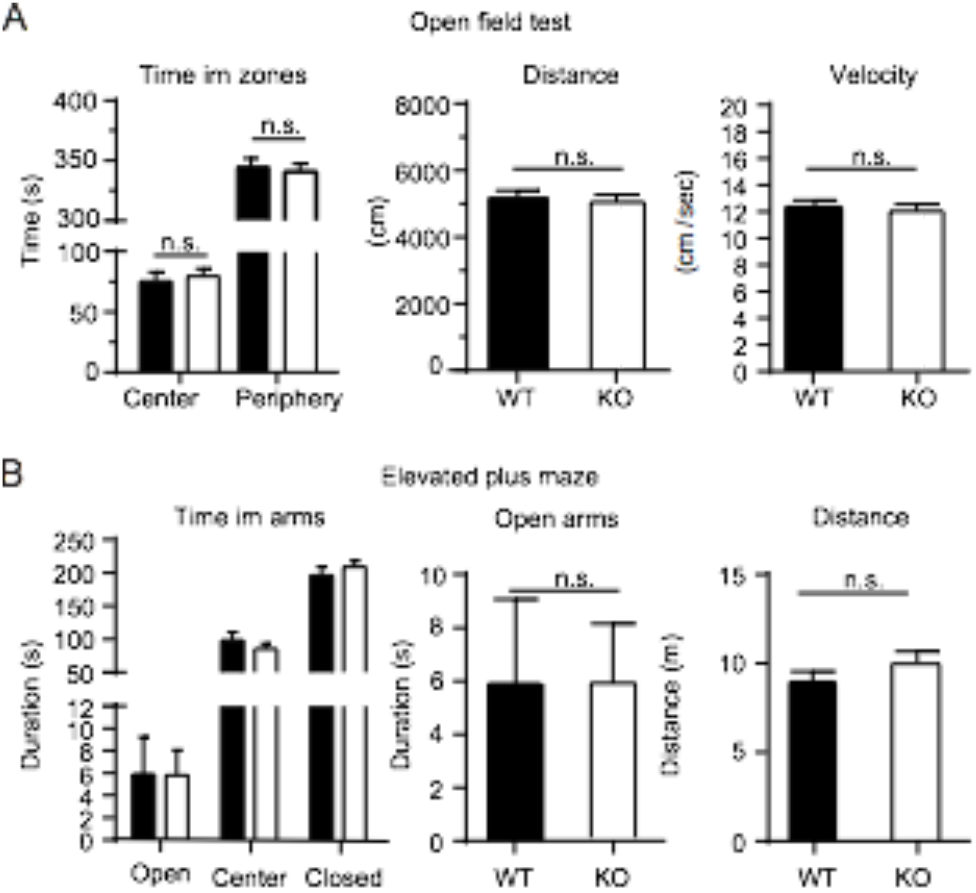
Unimpaired basic locomotion and no differences in anxiety as tested by the open field test and the elevated plus maze test.

**Supplemental Figure 4.**
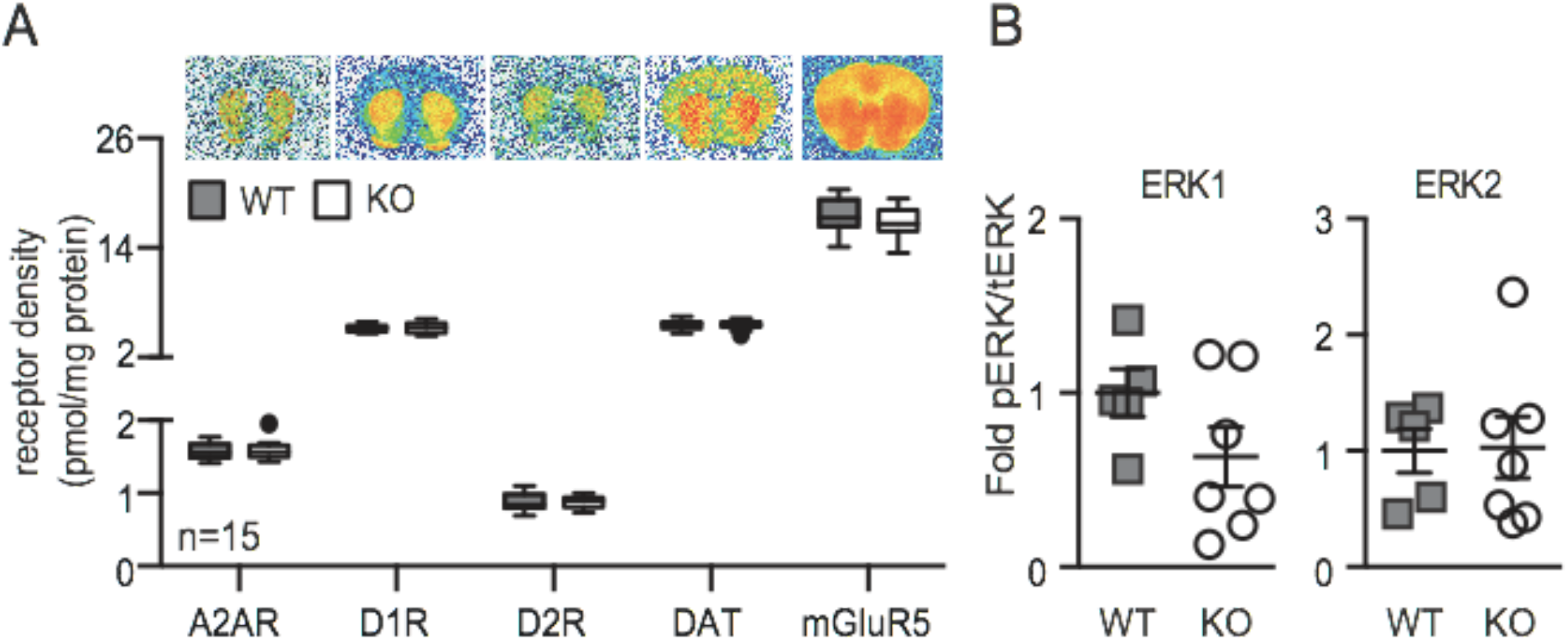
NECAB2 deficiency does not affect function or abundance of striatal GPCRs. **(A)** No differences in receptor density of striatal receptors A_2A_R, mGluR_5_, D1R, D2R and the dopamine transporter (DAT) in wildtype (WT) and *Necab2* knockout (KO) mice measured by receptor autoradiography. Data represent Tukey plots obtained from 15 WT and KO mice. **(C)** Similar ERK phosphorylation between WT and KO striatal tissue. Each data point represents the ratio of phosphorylated and total ERK from one mouse quantitated by immunoblotting.

